# Evaluating AI-Assisted Customer Verification for Synthetic Nucleic Acid Screening

**DOI:** 10.64898/2026.02.27.708645

**Authors:** Alejandro Acelas, Hanna Pálya, Kevin Flyangolts, Paul-Enguerrand Fady, Cassidy Nelson

**Affiliations:** The Centre for Long-Term Resilience, London, United Kingdom; University of Warwick, Coventry, United Kingdom; Aclid, New York, United States

**Keywords:** biosecurity, large language models, legitimacy screening, know your customer, sequence of concern, gene synthesis, human-in-the-loop, dual-use biotechnology

## Abstract

Legitimacy screening, the process of verifying the identity and purpose of customers ordering synthetic nucleic acids, is a primary safeguard against the misuse of synthetic biology. However, the associated costs discourage the adoption of screening practices. To evaluate whether AI tools can facilitate this process, we tested five large language models on five verification tasks using customer profiles of life sciences researchers from around the world. We compared AI performance against an expert human baseline on flag accuracy, source quality, source fidelity, and cost.

Flag accuracy of the best-performing model (Gemini 2.5 Pro with four bibliographic and sanctions APIs) was statistically indistinguishable from the human baseline at 90.2% and 89.0% (*n* = 41). Gemini 2.5 Pro performed at or above the human baseline on source quality and fidelity, at roughly one-tenth of the cost ($1.18 vs. $14.04 per customer). For information-gathering tasks, which excluded the human review step, costs averaged $0.23 per customer, around 50 times cheaper than human screening. These results support piloting AI assistance at the information-gathering step of legitimacy screening at providers of synthetic nucleic acids and other dual-use biotechnology products, with human reviewers retaining authority over follow-up communication and order fulfillment decisions.

## 1 Introduction

Synthetic nucleic acids are unique among commercially available laboratory reagents: they can be used directly to create pathogens through standard laboratory techniques, without requiring a live pathogen culture [Cello et al., 2002]. The overwhelming majority of synthetic nucleic acid use cases are beneficial—from vaccine development and diagnostics to industrial biotechnology applications—and legitimate research often requires access to sequences derived from dangerous pathogens [Tan et al., 2021]. However, a malicious actor could purchase commercially available synthetic nucleic acids to create dangerous pathogens [Haddad, 2025]. The sequences that could enable this misuse of synthetic nucleic acids are called “sequences of concern” (SOCs). Screening customers who order SOCs is a primary safeguard against misuse.

As nucleic acid synthesis becomes more accessible, strengthening these safeguards becomes more important. Organizations like the International Gene Synthesis Consortium (IGSC) have committed to two forms of screening. First, sequence screening compares the ordered sequence against databases of sequences from regulated pathogens. Second, legitimacy screening verifies that customers have legitimate reasons to order flagged sequences [International Gene Synthesis Consortium, 2024]. However, widespread adoption of sequence and legitimacy screening remains limited, with cost as a major barrier [Crawford et al., 2024]. For gene-length orders above 200 basepairs, recent work reports a per-order cost of £0.95 for legitimacy screening compared to £0.10 for sequence screening—almost an order of magnitude difference [Fady et al., 2025a]. When considering orders specifically flagged by sequence screening, the costs rise to £73 for legitimacy screening compared to £0.53 for sequence screening [Fady et al., 2025b].

Currently, no jurisdiction explicitly requires providers of synthetic nucleic acids to screen customers. While anti-terrorism laws create implicit obligations, these are neither standardized across countries nor grounded in a shared risk framework [Wheeler et al., 2024]. The EU Biotech Act may introduce customer screening requirements in late 2026, but the regulation is still in draft form and subject to change [European Commission, 2025]. Screening practices therefore vary substantially across providers and are currently in flux, creating uneven security guarantees against self-amplifying biological threats which easily cross borders.

Legitimacy screening typically involves two phases: onboarding and follow-up. Onboarding checks involve the verification of basic customer information such as name checks against watchlists, institutional affiliation, email domain ownership, and stated research purpose. When a SOC is flagged, follow-up screening determines whether the customer has a legitimate need for it. This phase requires gathering information from publications, patents, and institutional records, and requesting documentation from customers such as grant awards or biosafety approvals. The information-gathering step is time-consuming but largely mechanical; as it involves searching public records and synthesizing information, it is a natural target for AI assistance.

The aim of this study was to evaluate whether AI tools can accelerate information gathering as part of legitimacy screening. We evaluated five large language models with web search capabilities and with additional specialized tools on five bibliographic and sanctions verification tasks within follow-up screening (not onboarding or fulfillment). The verification tasks were: checking each customer’s institutional affiliation, institution type, and email domain ownership, as well as sanctions screening, and background work search. We found that AI-assisted screening reduced total per-customer legitimacy screening costs more than tenfold compared to manual screening ($1.18 versus $14.04), with accuracy comparable to our human baseline on the tasks evaluated. For tasks AI could perform without human intervention, costs averaged $0.23 per customer, which was around 50 times cheaper than human screening.

## 2 Methods

### 2.1 Task Definition

AI and human screeners were evaluated on five verification tasks relevant to customer follow-up screening:

1. **Institutional affiliation verification**: Confirm the customer is currently affiliated with their claimed institution.
2. **Institution type verification**: Confirm the institution is a legitimate life sciences research organization or company.
3. **Email domain verification**: Confirm the customer’s email domain belongs to their claimed institution.
4. **Sanctions screening**: Check whether the customer or their institution appears on export control or sanctions lists.
5. **Relevant work search**: Find publications, patents, or other work from the customer or their institution related to the ordered sequence.

For tasks 1–4, screeners made a choice from one of three outcomes: flag (concern identified or information not found), no flag (verified without concern), or undetermined (insufficient evidence to decide). For task 5, screeners reported relevant work without adding flag annotations.

These tasks were selected because they can often be resolved using publicly available information, they appear in existing screening guidance [Alexanian and Carter, 2024], and they represent the information-gathering phase of legitimacy screening rather than the judgment-intensive decision phase.

After information gathering, a human reviewer decided whether to ship the order, reject it, or request additional documentation from the customer. Screener performance was evaluated on the information-gathering tasks only. The order fulfillment decision was not evaluated, but its time and cost were included in our cost estimates.

### 2.2 Dataset

The primary dataset used for evaluating human and AI screeners had 41 customer profiles, each paired with a simulated order for a sequence of concern. Each profile included the name, institutional affiliation, email address of the customer, as well as a reference work (publication, patent, or news-reported activity) through which we identified the customer. To ensure data protection and prevent any false association between real researchers and simulated biosecurity threats, all personally identifiable information (PII) was strictly anonymised prior to the publication of the dataset.

Flag accuracy was manually graded on this dataset, while source quality, source fidelity, and work relevance were graded automatically with LLMs. AI models were also evaluated on an extended dataset of 134 customer profiles using automatic LLM-grading on source quality, source fidelity, and work relevance (see **Supplementary Material**, Section 7.2).

To simulate realistic screening scenarios, the customer dataset was composed of researchers identified through publications and patents. Some of these involved work on controlled pathogens, others on proteins that would not trigger screening concerns. Sampling was done across geographic regions and institutional types, and used the U.S. Consolidated Screening List (CSL) to include profiles that should trigger sanctions-related flags.

#### 2.2.1 Customer Types

Four groups of customer profiles were constructed:

**Academic users of SOCs:** Life science researchers from academic institutions with documented laboratory work on SOCs.

**Industry users of SOCs:** Researchers at life science companies with documented work on SOCs, performed either by the customer or by their institution.

**Customers in the U.S. Consolidated Screening List:** Researchers at academic institutions appearing on the CSL, with documented work on SOCs.

**General life science researchers:** Researchers from academia and industry with documented laboratory work on non-SOC sequences.

All profiles in the dataset—including general life science researchers identified through non-SOC work—were assigned simulated orders from a standardized list of 22 SOC proteins. The SOC proteins were selected such that it would be reasonable to order them for vaccine or therapeutics research (see **Supplementary Material**, Section 1).

Screeners therefore encountered cases where researchers appeared to order SOCs without relevant background—a mismatch they were expected to identify and note.

**Table 1:** Dataset composition by customer category. Numbers outside parentheses are the 41-profile primary dataset; numbers in parentheses are the extended 134-profile dataset. Region is determined by institution location. Institutional email indicates profiles with a business, academic, or government email domain.

|  | Academic<br>SOC Users | Industry<br>SOC Users | Life Sci.<br>Researchers | CSL<br>Academics | Total |
| --- | --- | --- | --- | --- | --- |
| Profiles | 16 (56) | 11 (24) | 7 (29) | 7 (25) | 41 (134) |
| **Region** |  |  |  |  |  |
| U.S. | 2 (6) | 4 (8) | 0 (1) | 0 (0) | 6 (15) |
| Europe + Australia | 5 (18) | 2 (6) | 0 (4) | 0 (6) | 7 (34) |
| China | 2 (8) | 3 (6) | 6 (19) | 1 (5) | 12 (38) |
| Other | 7 (24) | 2 (4) | 1 (5) | 6 (14) | 16 (47) |
| Inst. email | 15 (49) | 10 (22) | 5 (18) | 3 (19) | 33 (108) |

#### 2.2.2 Data Collection

Academic researchers were identified by searching for each SOC in Google Scholar. To promote geographic diversity, names sampled from a random name generator weighted toward common names in different world regions were included in some of the searches.

Publications that met three criteria were selected. First, they had to be published after 2021, to increase the likelihood that affiliations remain current. Second, they had to involve laboratory work on the searched protein, rather than purely computational or epidemiological analysis. Third, they had to plausibly require a synthetic nucleic acid. Results were filtered for authors with a publicly available email address or an institutional email domain identifiable through Google Scholar, ORCID, or similar research profiles.

Industry researchers were identified through two search strategies, one based on patents and the other based on news articles. For patent-based collection, PatentScope was queried using organism names, with patent holders from outside the US and EU being prioritized to encourage geographic diversity. Large pharmaceutical firms were excluded, with focus instead placed on smaller companies with staff rosters that are harder to verify and more likely to require individual legitimacy screening. For news-based collection, Google was searched for articles combining organism names with terms like “vaccine,” “diagnostics,” and “startup,” and researchers in laboratory roles at the resulting companies were then identified via LinkedIn.

Sanctioned researchers were identified through the manual review of the U.S. Consolidated Screening List for institutions conducting biological research.These were primarily academic institutions in China, Russia, and Iran. The institution names and the pathogens of SOCs were then searched in Google Scholar to find publications by authors at those institutions.

### 2.3 Screeners

#### 2.3.1 AI Models

Five large language models were tested, each with web search capabilities: Claude Sonnet 4 (Anthropic), Gemini 2.5 Pro (Google), Grok 4 (xAI), GLM 4.6 (Zhipu AI), and Minimax M2 (MiniMax). The latest available models were selected from major commercial providers released before November 1, 2025. GLM 4.6 and MiniMax M2 were included as competitive open-source alternatives based on public benchmark performance and LMArena rankings. Several models were excluded because they frequently refused to complete the screening task, citing biosecurity concerns, including OpenAI’s GPT-5, o1, and o3, and Anthropic’s Claude 4.5 Sonnet.

Each model was tested under two conditions:

**Web search only:** Models had access to web search via Tavily, a search API designed for AI applications, and no other tools.

**Web search and specialized tools:** Models had access to web search and four additional tools:

- *Consolidated Screening List API:* Searches the U.S. government’s consolidated list of sanctioned entities and restricted persons.
- *Europe PMC:* Searches Europe PubMed Central for scientific articles by author, institution, topic, or ORCID identifier.
- *ORCID profile:* Retrieves researcher profile information including affiliations, employment history, and recent publications.
- *ORCID works search:* Searches a researcher’s full ORCID publication list by keywords.

Both conditions used identical screening prompts (see **Supplementary Material**, Section 3).

#### 2.3.2 Human Baseline

Collecting customer profiles and grading outputs was conducted by the lead author, who did not serve as a screener.

Two coauthors served as the expert human baseline. One had spent over a year developing tools and standards for legitimacy screening, and the other works full-time building compliance software—including legitimacy screening tools—used by gene synthesis providers. Although neither received task-specific training, both could draw on this domain background, and on their familiarity with the research plan, to complete the task.

They were given an earlier version of the screening instructions (developed while optimizing prompts for model performance) and had not previously seen the profiles they evaluated. They submitted responses through a custom interface that enforced the same output format as model responses.

The interface recorded snapshots of each screener’s work at approximately 5 and 30 minutes. If they submitted a final answer earlier, that submission was used as the snapshot for that time point. Screeners were expected to complete each profile within 30 minutes, but there was no hard time cap; submissions after 30 minutes were accepted as received. Each evaluator screened 20 or 21 profiles, covering the full primary dataset. Profiles were randomly sampled with the constraint that no two shared the same reference work.

The 5-minute snapshot was collected to estimate human performance under a time budget comparable to AI models, which averaged under two minutes per profile. However, human responses at 5 minutes were typically incomplete, and this baseline underperformed AI models by a large margin. Unless otherwise noted, “human baseline” refers to the final submission throughout.

### 2.4 Evaluation Metrics

Each of the five verification tasks were evaluated on three binary metrics (whether the screener flagged correctly, and two response-quality metrics), yielding 15 measurements per customer profile. Grading all non-flag metrics manually was infeasible at this scale: the primary dataset alone required 5,412 such judgments, and the extended 134-profile dataset added a further 14,740. Gemini 2.5 Flash was therefore used in an LLM-as-a-judge setup for source quality, source fidelity, and work relevance, an established practice in LLM capability evaluations [Zheng et al., 2023, Chiang and yi Lee, 2023]. Flag accuracy, the most consequential binary outcome, was graded manually by comparing each screener’s flag choice against ground truth.

#### 2.4.1 Verification Tasks (Tasks 1–4)

##### Flag accuracy

Whether the response’s choice (flag, no flag, or undetermined) matches ground truth. The lead author established ground truth by manually reviewing all model and human responses, reading each screener’s justification and checking cited sources against the primary record. Unresolved cases were recorded as Ambiguous Case under the error taxonomy (see Section 2.5.1) and excluded from flag accuracy. None of the human-baseline screeners were involved in ground truth annotation; the annotator was not blind to screener identity, which risks systematic bias. Dual-annotator adjudication was not feasible given project resources, though the blind re-grading reported in Section (*κ* = 0.63) provides partial, indirect validation.

##### Source quality

Whether cited sources meet standards for independent verification. A source is acceptable if it exists independently of the customer and has editorial oversight. Acceptable sources include government registries, peer-reviewed publications, patent filings, regulatory submissions, business registrations, and established research profile aggregators. Unacceptable sources include LinkedIn profiles, personal websites, Wikipedia, and social media sites. A response passes this metric if all cited sources are acceptable; responses citing any unacceptable source, or no sources at all, fail.

##### Source fidelity

Whether factual claims are directly supported by cited sources. A claim fails if the cited source contradicts it, no cited source provides relevant information, or the claim presents speculation as established fact. Claims reporting absence of evidence (e.g., “no matching records found”) passed by default, as negative claims are inherently difficult to source. Empty responses automatically failed.

#### 2.4.2 Background Work Task (Task 5)

Background work was evaluated on source quality and fidelity using the criteria above, as well as a relevance metric.

##### Work relevance

Whether the response identifies work at least as relevant as the reference work used to include the customer in our dataset. Relevance is scored 0–5 based on three factors:

1. *Proximity to customer:* Work authored by the customer directly (higher) versus produced by their institution (lower).
2. *Organism proximity:* Same organism as the order (higher), closely related organism (middle), or unrelated (lower).
3. *Laboratory involvement:* Hands-on experimentation (higher) versus purely computational work (lower).

A response passed if at least one retrieved source scored at or above the relevance level of the reference work.

### 2.5 Prompt Development

The same task prompts were used across all models. They were developed through iterative refinement on ten customer profiles. After each iteration, the outputs were reviewed for common errors such as citing unrelated sources, making overly broad claims, failing to search multiple sources before concluding information was unavailable, and failing to comply with sequential instructions. Instructions were rewritten and consolidated multiple times to improve task adherence (see **Supplementary Material**, Section 3).

Evaluation prompts underwent a similar iterative process to improve alignment with human judgment. To validate the LLM-as-judge approach, the lead author independently re-graded a sample of 135 evaluation outputs without access to the original model grades. Agreement reached 83%, compared to 63% expected by chance (Cohen’s *κ* = 0.63). This falls within the “substantial agreement” range (0.61–0.80) on the Landis and Koch [1977] interpretation scale (see **Supplementary Material**, Section 4). Full blindness to the screener identity was not possible for either AI or human responses: model-specific output patterns (phrasing, response length) were recognizable to the re-grader from the earlier prompt-iteration work, and the two human screeners’ free-text justifications, although submitted through the same structured interface, differed in style and length from each other and from any model.

Due to the manual work required for flag accuracy grading, we only graded flag choices after settling on the final prompts provided to human and AI models. As a result, this may have underestimated flag accuracy performance for both AI and human screeners, as no iterative refinement was possible based on flag choice feedback.

#### 2.5.1 Flag Error Analysis

A random sample of 126 flag errors across AI and human screeners was reviewed and assigned each to one of five categories:

- **Criterion Deviation:** The screener’s flag choice conflicted with the criteria used to establish ground truth. These criteria were often specified explicitly in the screening prompt, though some recurring edge cases were not addressed directly and required interpretation.
- **Search Failure:** The screener did not locate information that was central to the flag choice but that other screeners successfully found.
- **Missing Response:** The screener produced no response in the relevant field, or the response was incomplete or not formatted according to task instructions.
- **Ambiguous Case:** The screener had access to all relevant information, but no clear rule in the flag choice criteria resolved the case, requiring a judgment call where reasonable screeners might disagree.
- **Source Misinterpretation:** The screener cited appropriate sources but misrepresented their content, leading to an incorrect flag.

#### 2.5.2 Cost and Time Estimation

Cost analysis was conducted using marginal costs, measuring the expense of screening each additional customer. Fixed costs such as prompt development, tool integration, and screener training can be amortized over a large number of cases and are excluded from our estimates. For both workflows, cost was measured throughout the initial screening step: from receiving a customer profile to recording an order decision. When the decision was to request follow-up, the cost of subsequent customer communication or verification of additional information was not included.

##### Human-only screening costs

Costs were estimated based on time required and hourly wage for comparable roles. Using advertised salaries for customer service positions at large DNA synthesis providers [Glassdoor, 2025], an hour of human screening was estimated at approximately $54. This figure depends heavily on the country in which a screener is employed. Due to limited salary data, it was treated as representative of industry-wide costs.

##### AI-aided screening costs

For information gathering, costs were based on each provider’s per-token API pricing (with input and output tokens priced separately) and the cost of each web search query requested by the model. For order fulfillment decisions, the human review process was timed and its costs were estimated using the same hourly rate.

For AI screening only, we also computed correlations between per-customer costs (excluding human review) and overall performance across model configurations to understand the possible trade-off between cost and accuracy.

## 3 Results

### 3.1 Pass Rates by Screener

As Figure 2 shows, AI models outperformed the human baseline on most metrics. On every metric other than flag accuracy, all models achieved a higher pass rate than the human screeners; the advantage reached statistical significance on claim support (3 of 10 configurations) and work relevance (4 of 10), but not on source reliability, where intervals overlap the human baseline across all configurations (**Supplementary Table S2**). Even for flag accuracy, the human baseline outperformed only two of the ten model configurations. The 5-minute-limit human baseline performed much worse than any other screener setup, largely because responses were practically always incomplete, leading to automatic failures on evaluation metrics.

**Figure 1:**
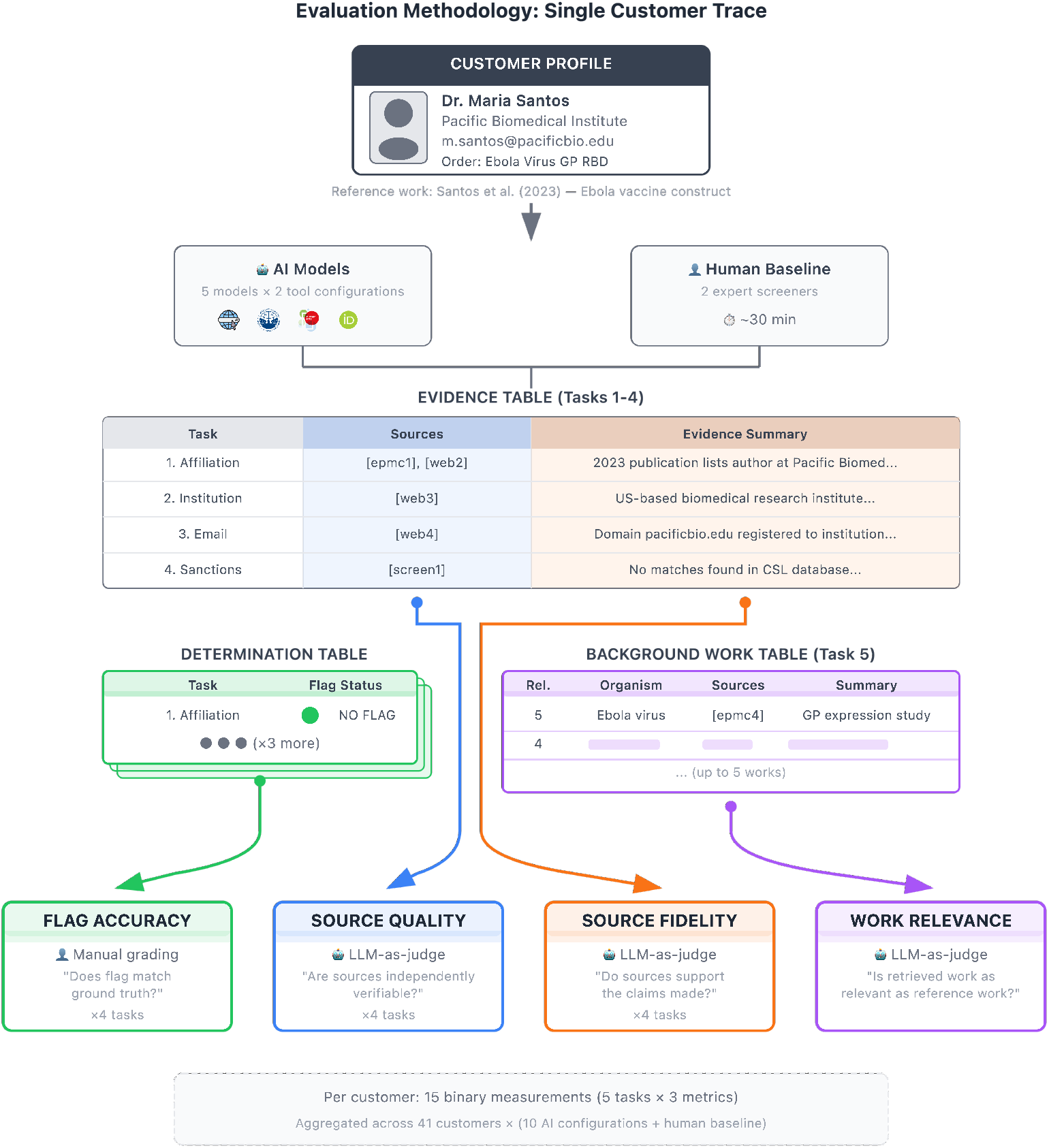
Evaluation methodology for AI-assisted legitimacy screening. Each customer profile includes a simulated order for a sequence of concern and a reference work (publication, patent or news article). Screeners produce structured outputs evaluated on three binary metrics per task: source quality, source fidelity, and flag accuracy / work relevance. The example shown uses a fictional customer profile for illustration purposes.

**Figure 2:**
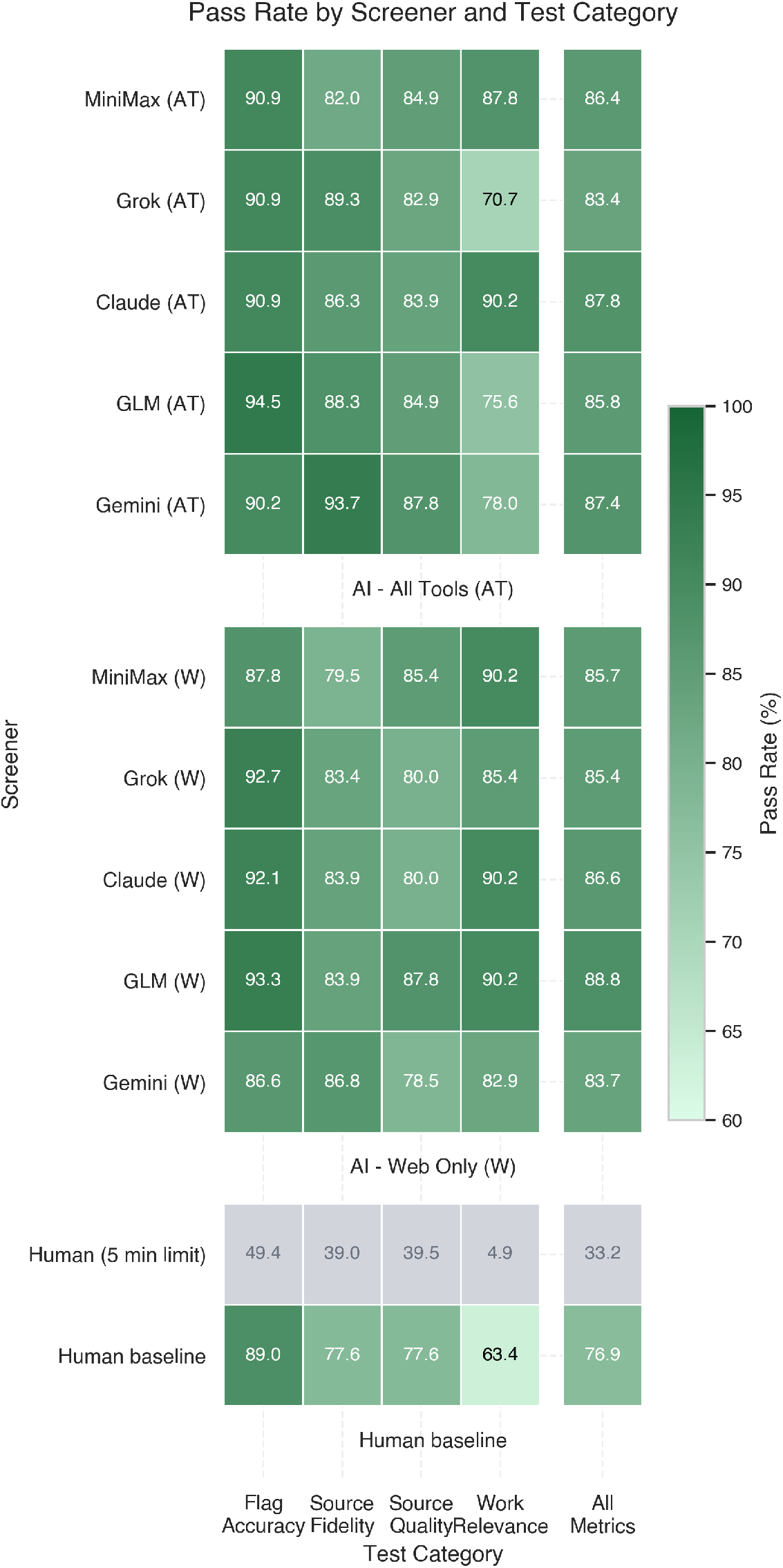
Pass rates by screener and metric. Each cell shows the percentage of evaluation tests passed across 41 customer profiles. Wilson 95% confidence intervals are given in **Supplementary Table S2**, and **Supplementary Figure S1** shows a forest plot of the row-wise overall rates. Columns show four metrics: flag accuracy (correct flag/no-flag/undetermined choice), source fidelity (claims supported by cited sources), source quality (sources are independently verified), and work relevance (work outputs listed are at least as relevant as a known reference).

**Figure 3:**
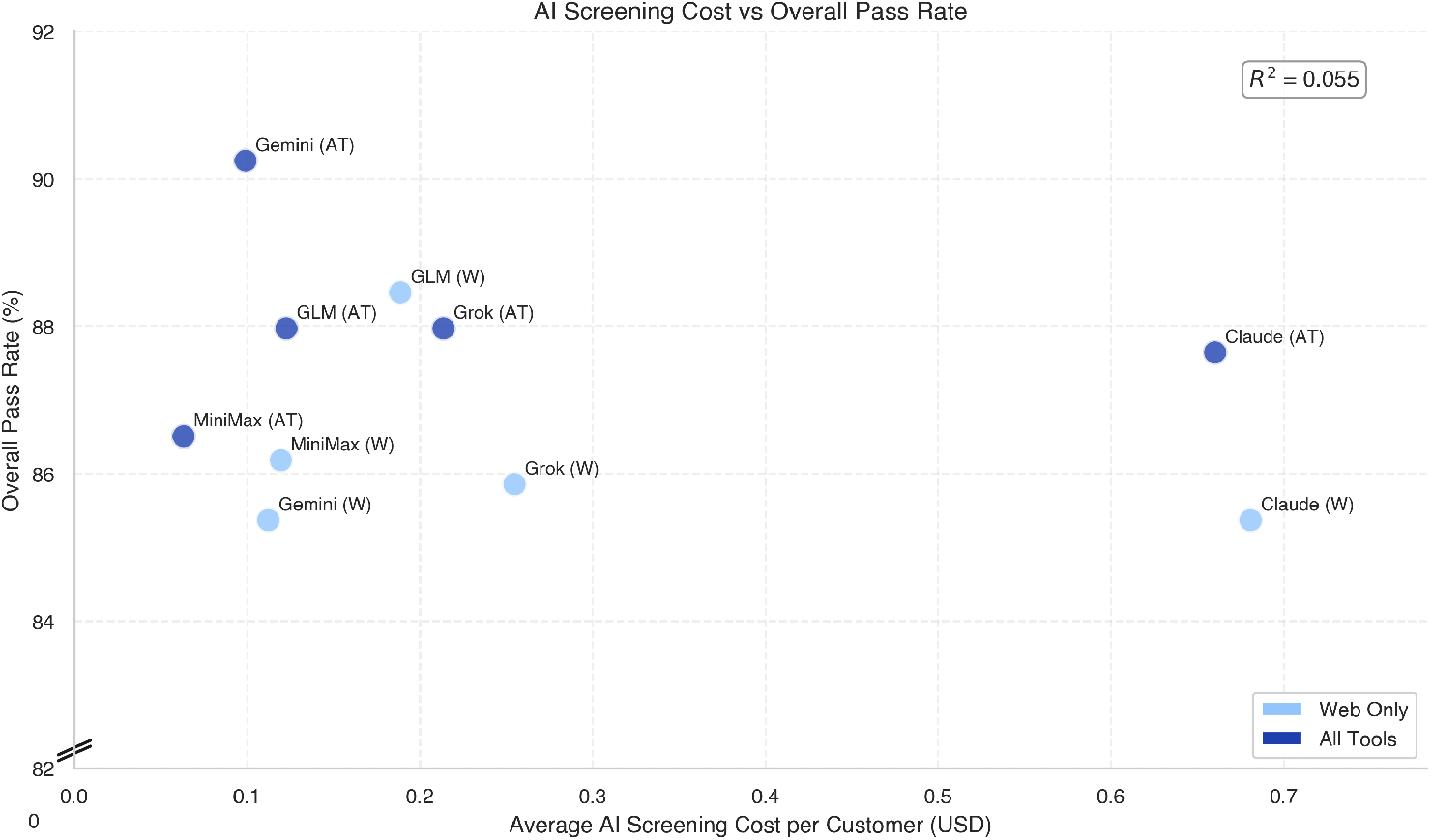
AI screening cost versus overall pass rate. Each point represents one model configuration (five models × two tool conditions). The x-axis shows average AI information-gathering cost per customer (excluding human review); the y-axis shows the overall pass rate across all four metrics (note the broken y-axis starting at approximately 83%). All values are averages across 41 customer profiles.

**Figure 4:**
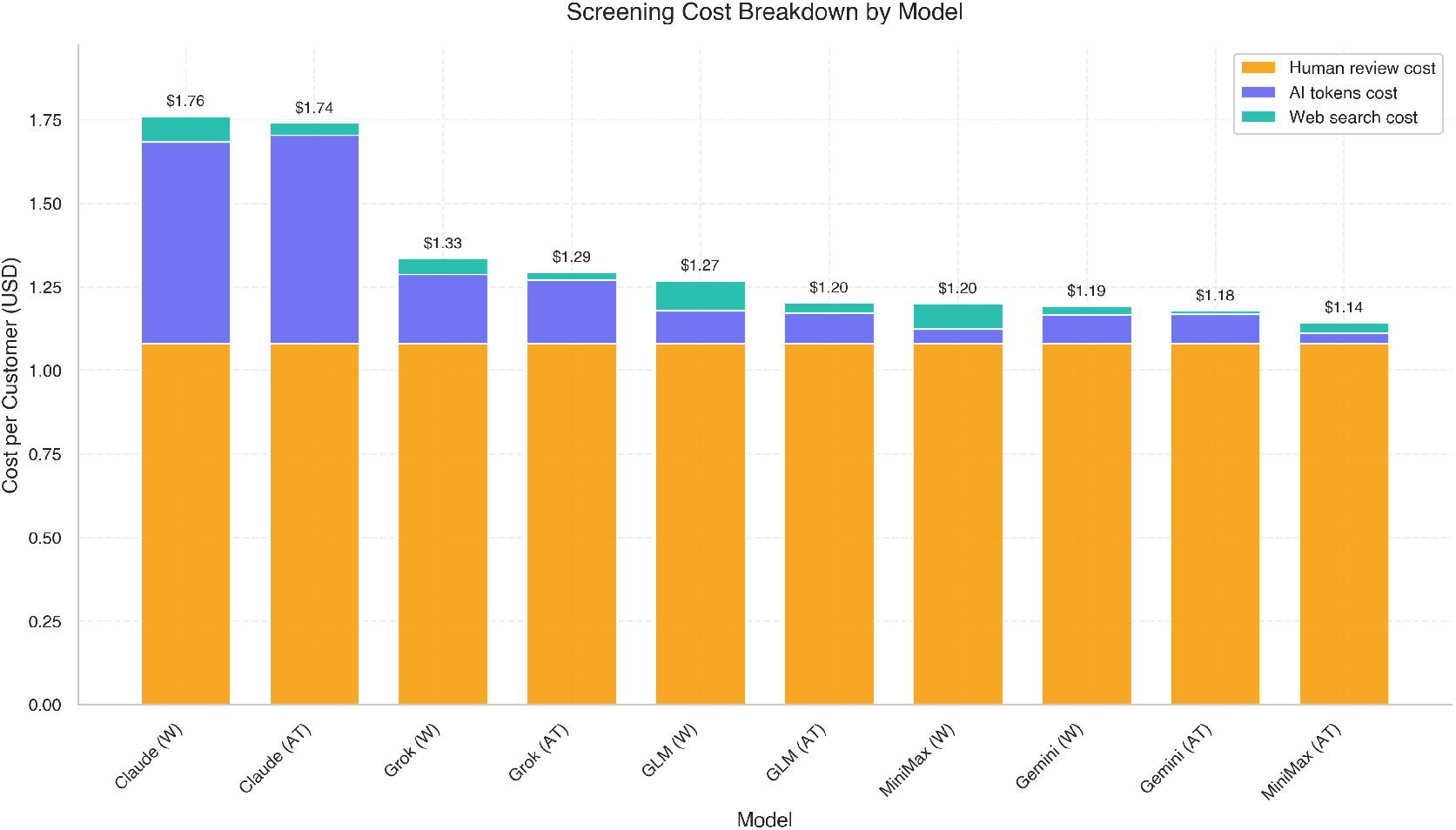
Per-customer screening cost breakdown by model configuration. Stacked bars show three cost components: human review of the AI-generated report (orange), AI token costs for input and output (purple), and web search queries via Tavily (teal). Human review cost was estimated only for the best performing model configuration (Gemini 2.5 Pro with all tools), and it is taken as constant across configurations (approximately $1.08). All values are averages across 41 customer profiles.

Across AI models, pass rates spanned a narrow range: the best-performing configuration (Gemini 2.5 Pro with specialized tools) passed 89.8% of tests, while the worst-performing (Gemini 2.5 Pro with web search only) passed 83.7%. Notably, this was the same underlying model in both cases, differing only in tool access.

Access to specialized tools (Consolidated Screening List API, Europe PMC, ORCID) affected performance differently depending on the task. For verification tasks (Tasks 1–4), tool access produced a modest but consistent improvement across all metrics. The improvement was most pronounced for sanctions screening (**Supplementary Figure J5**), where the Consolidated Screening List API provided direct access to authoritative data otherwise available only through downloadable files or specialized interfaces.

For work relevance (Task 5), the pattern reversed: models with tool access scored lower than those with web search only. The drop was concentrated among industry customers (**Supplementary Figure E4**), while academic and other customer types showed little difference between configurations.

This likely reflects the fact that models with tool access performed notably fewer web searches (**Supplementary Figure J1**), making them more likely to miss patents or news articles—sources not indexed by Europe PMC or ORCID, which cover primarily academic publications. For most profiles in our dataset, this substitution had little effect, as publication databases contained comparable evidence. However, industry customers were specifically selected based on non-publication work outputs, so tool access steered models away from the most relevant sources for these customers.

### 3.2 Cost Comparison

All AI models produced customer reports faster and more cheaply than either human baseline. Comparing total costs—including human review of AI-generated reports—the most expensive model tested (Claude Sonnet 4, $1.74 total) was approximately 8 times cheaper than fully manual screening ($14.04), while the cheapest configuration (MiniMax M2 with tools, $1.14 total) was approximately 12 times cheaper. The cost reductions were more dramatic when comparing information gathering alone: AI models averaged $0.23–$0.27 per customer, roughly 50–60 times cheaper than the human baseline.

Across AI models, cost per customer did not correlate with overall pass rates (*R*^2^ = 0.01). The best-performing model, Gemini 2.5 Pro with all tools, was also the second cheapest configuration. Although the open-source models (GLM 4.6 and MiniMax M2) had per-token prices 2–8*×* lower than Gemini 2.5 Pro, additional web search queries and higher token consumption (often from extra tool calls) eliminated any meaningful cost advantage.

**Table 2:** Per-customer screening costs and processing times. “Information gathering” covers Tasks 1–5 only; “total cost” adds the time cost of human review of the AI-generated report. For human baselines, these phases were not separated, so only totals are reported. Human costs estimated at $54/hour based on advertised salaries at a large DNA synthesis provider. AI costs include per-token API pricing and Tavily web search queries ($0.08/query); other tools were cost-free. All figures are averages across 41 customer profiles.

| Model | Info. Gathering<br>Cost | Total<br>Cost | Info. Gathering<br>Time | Total<br>Time |
| --- | --- | --- | --- | --- |
| *Human baseline* |  |  |  |  |
| Human (5 min limit) | — | \$5.43 | — | 6.0 min |
| Human baseline | — | \$14.04 | — | 15.6 min |
| *AI models (all tools)* |  |  |  |  |
| Claude Sonnet 4 | \$0.66 | \$1.74 | 1.5 min | 2.7 min |
| Grok 4 | \$0.21 | \$1.29 | 2.6 min | 3.8 min |
| Gemini 2.5 Pro | \$0.10 | \$1.18 | 1.3 min | 2.5 min |
| GLM 4.6 | \$0.12 | \$1.20 | 1.7 min | 2.9 min |
| MiniMax M2 | \$0.06 | \$1.14 | 2.2 min | 3.4 min |
| *Average (all tools)* | *\$0.23* | *\$1.31* | *1.9 min* | *3.1 min* |
| *AI models (web-only)* |  |  |  |  |
| Claude Sonnet 4 | \$0.68 | \$1.76 | 1.0 min | 2.2 min |
| Grok 4 | \$0.26 | \$1.33 | 2.7 min | 3.9 min |
| Gemini 2.5 Pro | \$0.11 | \$1.19 | 1.3 min | 2.5 min |
| GLM 4.6 | \$0.19 | \$1.27 | 1.4 min | 2.6 min |
| MiniMax M2 | \$0.12 | \$1.20 | 2.5 min | 3.7 min |
| *Average (web-only)* | *\$0.27* | *\$1.35* | *1.8 min* | *3.0 min* |

As web search accounted for 60% of total processing cost of using MiniMax M2, alternative web search providers—including custom tools offered by some model developers—could reduce costs further. However, the lack of standardization across such tools would require adapting prompts and source extraction for each provider.

### 3.3 Flag Error Analysis

#### 3.3.1 Error Distribution

Except for errors in the search failure category, the frequency of each error type was similar across screeners. Table 3 summarizes error category distributions across screener types. Criterion deviation was the most common category, accounting for 39.7% of all errors. On inspection, a small number of recurring patterns accounted for the majority of mistakes in this category, such as failing to flag personal (non-institutional) email addresses used by otherwise reputable researchers in China.

**Table 3:** Distribution of flag errors by category and screener type. Percentages are computed within each screener type (column), showing what share of that screener type’s errors fell into each category. Counts are in parentheses; the 126 total errors were sampled across all screeners. “Human Baseline” combines both the 5-minute-limit and final-submission snapshots. Examples illustrate representative instances; error categories are defined in the text.

| Error Category | Example | Human Baseline | AI (All Tools) | AI (Web Only) | Overall |
| --- | --- | --- | --- | --- | --- |
| Criterion Deviation | Flagging a biology dept. as “not in life sciences” because its university appears on a sanctions list | 35% (6) | 39% (19) | 42% (25) | 40% (50) |
| Search Failure | Missing that a researcher moved to a different institution | 35% (6) | 35% (17) | 17% (10) | 26% (33) |
| Missing Response | Empty response field or abruptly terminated output | 18% (3) | 20% (10) | 23% (14) | 21% (27) |
| Ambiguous Case | Email domain matches recently outdated company domain | 12% (2) | 6% (3) | 15% (9) | 11% (14) |
| Source Misinterp. | Assuming connections between nearby organizations without evidence | 0% (0) | 0% (0) | 2% (1) | 1% (1) |

We expect that including explicit guidance addressing these recurring cases could substantially reduce criterion deviation errors for both AI and human screeners.

Search failure was the next most common error type. Notably, web-only models had a lower rate (16.7%) than AI models with access to specialized tools or human screeners. Although we did not examine this pattern in detail, it resembles what we observed for work relevance: models with additional tools performed fewer web searches, potentially missing information available only through general web results.

Missing responses accounted for 21.4% of errors. For human screeners, these typically arose from deliberate decisions to skip a criterion—for example, skipping remaining checks after confirming a customer’s institution appears on a sanctions list—that coincided with a flag error. For AI models, missing responses often resulted from outputs that did not adhere to the requested table format or, occasionally, from extraction failures when parsing model responses.

Ambiguous cases and source misinterpretation were the least common, accounting for 11.1% and 0.8% of errors respectively. Ambiguous cases typically involved conflicting or partial evidence that made it difficult to establish clear ground truth. Source misinterpretation was rare despite the fact that around 15% of screener responses contained some factual inaccuracy, as measured by the source fidelity metric. This suggests that most of these inaccuracies were minor and did not affect the final flag choice.

#### 3.3.2 Geographic Variation

Error rates varied by customer region (Figure 5). European customers had the highest pass rates, potentially reflecting better documentation in the sources most available to screeners (English-language web search results and EuropePMC publications). Chinese customers showed higher missed flag rates on domain verification—15.0% compared to under 5% for other regions—as screeners often did not flag otherwise reputable researchers who used personal (non-institutional) email addresses.

**Figure 5:**
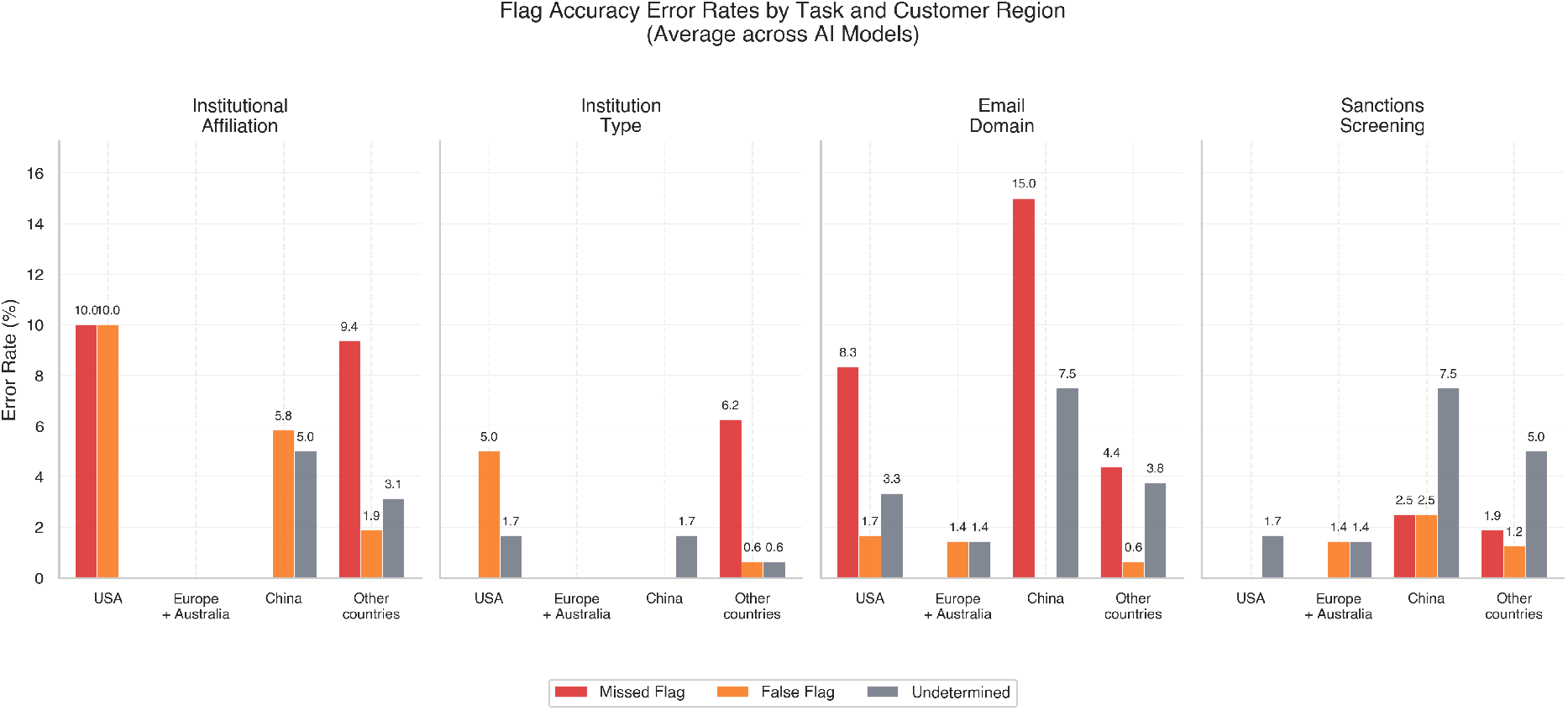
Flag accuracy error rates by task and customer region, averaged across all AI model configurations. Each panel shows one verification task; bars indicate the rate of missed flags (red; screener reported no flag when ground truth was flag), false flags (orange; screener reported flag when ground truth was no flag), and undetermined responses (gray; screener reported undetermined when ground truth was either flag or no flag). Regions are defined by institution location: U.S. (6 profiles), Europe + Australia (7 profiles), China (12 profiles), and Other countries (16 profiles). Error rates are computed as a percentage of all screener–profile pairs within each region.

Chinese customers also showed elevated undetermined rates on sanctions screening (7.5%), driven by institutions that appear on some countries’ sanctions lists but not others, creating genuine ambiguity about whether a flag is warranted. U.S. customers showed unexpectedly high affiliation verification errors (10.0% missed flags, 10.0% false flags), but we did not identify any consistent pattern when reviewing the full response transcripts, and the small sample size (6 profiles) limits the conclusions that can be drawn from this region.

## 4 Discussion

### 4.1 Key Findings

We evaluated AI models on the information-gathering step of follow-up screening of customers who ordered sequences of concern, comparing models with web search alone against those with additional specialized tools (Consolidated Screening List API, Europe PMC, ORCID), and against a human baseline. While our evaluation took place outside a nucleic acid synthesis company, we simulated realistic screening scenarios with customers who have verifiable research backgrounds. These results warrant piloting AI assistance at the information-gathering step of legitimacy screening at DNA synthesis companies and possibly other providers with similar risk profiles (e.g., cloud labs and pathogen repositories). This aligns with existing screening guidelines that pair sequence screening with legitimacy screening and recommend follow-up when orders are flagged [U.S. Department of Health and Human Services, 2023, International Gene Synthesis Consortium, 2024].

AI models matched or exceeded the human baseline on most metrics, with the best model (Gemini 2.5 Pro with tools) achieving an 89.8% average pass rate compared to 79.7% for humans. However, we invested substantially more effort in prompt development than in training human screeners. The human screeners were briefed on the criteria but not the precise rubrics, and unlike the prompts, their responses were never iterated against rubric scores. Experienced screeners with domain expertise might outperform current models on ambiguous cases requiring contextual judgment—cases our evaluation included but may have undersampled. This performance pattern reflects how LLMs are trained: instruction-tuned models excel at following written procedures and structured checklists, but they do not learn from experience during deployment unless explicitly updated through fine-tuning. Our approach, encoding screening guidance in detailed prompts with explicit rules for acceptable sources, plays to these strengths while avoiding the need for per-case adaptation.

The same prompt-driven design exposes a distinctive risk. Large language models are susceptible to prompt injection, in which instructions embedded in retrieved web pages, submitted documents, or customer-supplied free-text fields override the screening prompt and steer the model toward a cleared determination [Greshake et al., 2023, Zhan et al., 2024]. Although reports of successful prompt injection attacks in the wild are currently rare, it is conceivable that they might become more common in the near future. In such cases, production deployment would need defenses such as restricting customer-controlled text to clearly delimited fields, separating retrieval from instruction-following, and validating flag decisions against tool-verified claims rather than the free-form response.

Models showed greater variation across source quality metrics than on flag accuracy. Flag accuracy pass rates clustered within a narrow range for both AI and human screeners. However, source quality and fidelity metrics showed wider spreads: Claude Sonnet 4 struggled with institution type verification sources where the human baseline excelled; conversely, Grok 4 with tools substantially outperformed humans on background work relevance.

AI models completed screening in approximately 3.1 minutes on average (including human review) compared to 15.6 minutes for the human baseline—roughly a 5-fold throughput advantage. This speed becomes increasingly important as the synthetic nucleic acid market grows at a projected compound annual growth rate of approximately 15% [Fady et al., 2025c]. AI-assisted screening enables providers to scale verification capacity without proportional increases in human labor.

Access to specialized tools provided modest but consistent improvements across most tasks. The advantage was most pronounced for sanctions screening, where models with Consolidated Screening List API access substantially outperformed web-only configurations. Sanctions lists are typically available only through APIs or downloadable files, not as indexed webpages, making direct API access particularly valuable. However, tool access reduced performance on background work search by steering models away from web searches that would find patents and news articles not indexed in academic databases. Our implementation queried only the U.S. Consolidated Screening List; integrating additional sanctions databases (EU lists, OFAC, UN Security Council) could further improve coverage. The applicable lists are ultimately the synthetic nucleic acid provider’s decision, informed by the law of its jurisdiction.

### 4.2 Implications for Screening Practice

AI-assisted legitimacy screening has attracted interest from multiple commercial actors, including biosecurity-focused startups [Aclid, 2026, TwentyTwo, 2026] and established synthesis providers. Several well-known providers have indicated they are exploring integration of such tools into their order systems (personal communications with industry representatives, 2025). AI assistance at the information-gathering step may become an established part of customer screening standards, particularly if recognized by regulatory and compliance frameworks.

Cost reduction is the primary driver for its adoption. The best-performing model (Gemini 2.5 Pro with tools) cost $1.18 per customer including human review, compared to $14.04 for manual screening—roughly a 10-fold reduction. Information-gathering tasks alone cost $0.23 per customer on average for AI models, roughly 50 times cheaper than the human baseline. Open-source tools for legitimacy screening [Acelas, 2026] could accelerate adoption by lowering barriers for providers who already screen and reducing costs for those considering implementation.

Crucially, the ship/follow-up/reject decision should remain with humans; fully autonomous AI screening risks introducing biases from language models to order decisions, and should be avoided until we have stronger performance guarantees. AI error rates, while lower than our human baseline on these information-gathering tasks, remain non-negligible for security-critical applications where false approvals could enable misuse or prevent legitimate research. Ideally, future standards would specify acceptable false-negative and false-positive rates consistent with the security goals of customer screening. Beyond meeting current screening standards, this human-in-the-loop design also lets the workflow adapt as new misuse strategies and information sources emerge, flagging cases for human judgment when the AI’s evidence is thin or inconsistent.

Framed this way, AI-assisted legitimacy screening is an adaptive verification layer rather than a fixed compliance checklist. The AI component can be re-prompted, re-tooled, or substituted as misuse vectors and information sources change; final authority and case-by-case learning stay with the human reviewer. This is the architecture implicit in current and forthcoming screening regimes, which assume a screener who updates procedure in response to new threats rather than one who executes a static rule [Wheeler et al., 2024, European Commission, 2025].

The tasks evaluated here could plausibly be part of one legitimacy screening workflow, but they could also be split into onboarding tasks and follow-up tasks.

Onboarding tasks typically include name checks against watchlists, verification of institutional affiliation and email domain, and confirmation of research purpose. Follow-up tasks, triggered when orders contain sequences of concern, aim to collect information about whether the customer has a legitimate need for the sequence. Follow-up often requires requesting additional documentation from customers (biosafety approvals, grant awards, publications) and then verifying it. AI excels at the verification component but is yet to be tested on tasks requiring human interaction to request additional information.

AI-assisted legitimacy screening sits within a broader ecosystem of measures for enhancing biosecurity in synthetic nucleic acid access. It can be strengthened by frontloading information collection. Standardized templates, such as those developed by IBBIS for customers ordering sequences of concern [International Biosecurity and Biosafety Initiative for Science, 2025], prompt customers to provide verification materials upfront. Whitelisting verified customers for specific sequences with periodic re-verification [Crawford et al., 2024] similarly converts requests into verification tasks. At present, the value of AI-assisted screening lies in accelerating these verification tasks that currently bottleneck workflows, while humans retain responsibility for requesting additional information and making final fulfillment decisions.

Emerging EU regulation is supportive of this approach. The proposed European Biotech Act [European Commission, 2025], published in December 2025, introduces harmonized EU-wide biosecurity obligations for “biotechnology products of concern,” requiring providers to assess the legitimate need of prospective customers through screening procedures explicitly likened to know-your-customer assessments. We believe that the integration of AI tools into KYC checks as described here would not classify as a high-risk AI system under the EU AI Act [European Parliament and Council of the European Union, 2024]. Providing synthetic nucleic acids does not fall within any of the Annex III high-risk categories. Further, the system is explicitly designed as human-in-the-loop: the final ship, follow-up, or reject decision remains with human operators, meaning the AI does not autonomously determine outcomes that affect individuals’ rights. Either condition alone would suffice to exempt the system from high-risk classification under Article 6.

Performance on realistic but non-deceptive profiles does not speak to behavior under deliberate deception. In a small adversarial case study adapted from a fabricated profile developed by the Engineering Biology Research Consortium, all ten configurations correctly caught most of the red flags, but 8 of 10 were fooled by a look-alike institution: the claimed “PathoSense Labs” at pathosense.org mimicked a legitimate Belgian biotech at pathosense.com, and most models imported accurate information about the real entity to clear the fabricated one (see **Supplementary Material**, Section 5). More systematic red-teaming on a more representative customer sample was out of scope here due to resource constraints, but some of the authors are tentatively planning such a follow-up.

### 4.3 Limitations

Our screening approach depends on publicly available information about the customer. This affects both AI and human screeners and is inherent to any information-gathering system. However, dependence on public documentation may introduce bias: well-documented customers receive faster verification, while customers with limited online presence—including researchers at institutions with minimal web visibility, early-career personnel, and stealth startups—face longer review times or heightened scrutiny. This could disproportionately affect researchers from low- and middle-income countries, who already face barriers to accessing biotechnology.

Selection bias in our dataset construction may inflate performance estimates for both AI and human screeners: every profile required discoverable publications, patents, or institutional affiliations—information needed both to construct realistic profiles and to let screeners verify them. This overrepresents established researchers relative to the typical DNA synthesis customer base, which also includes laboratory technicians, students, and industry personnel with minimal public records. Performance on our benchmark may therefore overestimate real-world accuracy for these less-documented customers, who will remain comparatively more reliant on direct communication and manual review. For similar reasons, our dataset may underrepresent the most ambiguous, judgment-intensive cases where experienced screeners’ contextual interpretation matters most, which could inflate AI performance compared to the human baseline.

Our primary dataset of 41 profiles also limits statistical power: at *α* = 0.05 and 80% power a paired McNemar test can only reliably detect flag-accuracy differences of roughly 10–14 percentage points between two screeners, though the tighter intervals on source quality, source fidelity, and work relevance (**Supplementary Table S2**) already identify metrics on which the best AI configurations significantly outperform the human baseline.

Our human baseline has three features that limit direct comparison with AI: it comprised two coauthor evaluators rather than professionally trained screeners; those evaluators received no task-specific training beyond an earlier version of the screening instructions; and the evaluation tasks were selected partly for their fit with AI strengths— structured tasks with publicly verifiable answers. Trained screeners with domain expertise would likely outperform this baseline, particularly on ambiguous cases requiring contextual judgment. In-situ evaluation at nucleic acid synthesis providers, comparing AI-assisted screening against trained screeners on their own customer flows, would be advisable before production deployment. Ground-truth grading could not be fully blinded either: although the submission interface enforced an identical output format for all screeners, model phrasing and the human screeners’ free-text justifications still made provenance partly recognizable to the annotator, leaving some risk of systematic bias.

## 5 Conclusion

This work builds on years of guidance on DNA synthesis legitimacy screening [Carter and Friedman, 2015, Alexanian and Carter, 2024, International Gene Synthesis Consortium, 2024]. We evaluated how commercially available and open-source LLMs perform on the information-gathering tasks of follow-up legitimacy screening, as outlined by Alexanian and Carter [2024] and present in various screening guidances. We demonstrated that AI assistance at this step is substantially less costly and faster than human-only screening, with accuracy comparable to our human baseline on the information-gathering tasks evaluated here.

These results support leveraging AI assistance at the information-gathering step of legitimacy screening as part of industry-wide biosecurity. Although we conducted this study with the DNA synthesis industry in mind, our results suggest the same approach could extend to legitimacy screening for any dual-use product in the bioeconomy. This includes physical assets like live viral or microbial cells and benchtop DNA synthesis equipment. Crucially, it also extends to digital assets, providing a much-needed scalable verification mechanism for gating access to frontier and specialist AI models with biological capabilities. Future work should explore hybrid AI–human systems where models flag uncertain cases for human review, tailored to provider-specific data, and integrated with broader customer onboarding workflows.

## Data Availability Statement

The materials to replicate the main results of this work are available on GitHub at https://github.com/alejoacelas/ai-kyc-dna-synthesis-frontiers. Artefacts derived from real customer information (full screener responses and ungraded outputs) are excluded from the public release to protect personal data; this, together with reliance on proprietary LLMs whose versions evolve and exhibit stochastic outputs, limits exact replication. The published pipeline, prompts, and scoring criteria are detailed enough for other teams to construct comparable datasets and evaluations.

## Author Contributions

AA, HP, KF, PEF, and CN contributed to the conceptualization of the study. AA, HP, and KF developed the methodology. AA developed the software, conducted the formal analysis, performed the investigation, provided resources, curated the data, and created the visualizations. HP and AA wrote the original draft. HP, KF, PEF, and CN reviewed and edited the manuscript. PEF and CN provided supervision and acquired funding. CN administered the project. All authors reviewed and approved the final manuscript.

## Acknowledgments

We thank Becky Mackelprang at the Engineering Biology Research Consortium (EBRC) for providing the adversarial case study profile used in our evaluation, and Janika Schmitt and Tessa Alexanian for helpful comments on earlier drafts of this manuscript. We also thank Frankie Di-Nozzi for project management and operations support. Acknowledgment does not imply endorsement of the paper or its findings.

AI assistants (Claude, Anthropic; Gemini, Google) were used to proofread and suggest improvements to the manuscript text, write code for the evaluation pipeline, data analysis, figure generation, and the human evaluation interface, and to format the LATEX submission.

## Supplementary Material

Supplementary material is provided as a separate document.

