## Supplementary Material for "Evaluating AI-Assisted Customer Verification for Synthetic Nucleic Acid Screening"

#### 1 Sequences of concern used in evaluation

All customer profiles in our dataset were assigned simulated orders from the following list of 22 proteins. These sequences are derived from controlled pathogens and would typically trigger follow-up screening at DNA synthesis providers. We selected sequences that would be plausible to order for legitimate vaccine or therapeutics research.

The "General life science researchers" customer category was identified through documented laboratory work on non-SOC sequences including antibodies (scFv fragments), CAR-T constructs, protein engineering targets, and diagnostic antigens.

- Ebola Virus Glycoprotein (GP) RBD
- Foot-and-mouth disease virus VP1
- H5N1 Hemagglutinin (HA) receptor-binding site
- H7N9 influenza HA receptor binding site
- Human T-lymphotropic virus (HTLV) Tax protein
- Human metapneumovirus attachment protein
- Kaposi's sarcoma-associated herpesvirus K1
- MERS-CoV spike RBD
- Measles virus fusion protein
- Merkel cell polyomavirus large T antigen
- Mumps virus small hydrophobic protein
- Newcastle disease virus fusion protein cleavage site
- Oropouche virus nucleocapsid
- Parainfluenza virus 3 hemagglutinin-neuraminidase

- Peste des petits ruminants virus fusion protein
- Respiratory syncytial virus fusion protein
- Rinderpest virus hemagglutinin
- SARS-CoV-2 Receptor Binding Domain (RBD)
- SFTS virus nucleocapsid protein
- Schmallenberg virus nucleocapsid
- Usutu virus envelope protein
- Zika virus NS1 protein

### 2 Statistical power and confidence intervals

#### 2.1 Per-screener pass rates with 95% confidence intervals

Figure 1 plots each screener's overall pass rate with its 95% Wilson score confidence interval ($n = 615$ binary outcomes per screener: 41 customer profiles $\times$ 15 measurements). The ten AI intervals overlap one another and overlap the 30-minute human baseline at 79.7% [76.3%, 82.7%]; only the 5-minute human baseline is clearly separated. Table 2 reports the same intervals broken out by metric, corresponding cell-for-cell to Figure 2. A paired McNemar exact test of best-AI (Gemini 2.5 Pro with all tools) against the 30-minute human baseline on flag accuracy returned $p = 1.000$ at the profile level (41 paired profiles, 7 vs. 8 discordant pairs) and $p = 0.824$ at the criterion level (164 items, 9 vs. 11 discordant pairs). We cannot distinguish the two screeners on this metric; at $n = 41$, the study has 80% power to detect differences of roughly 10–14 percentage points depending on discordance rate (Table 4).

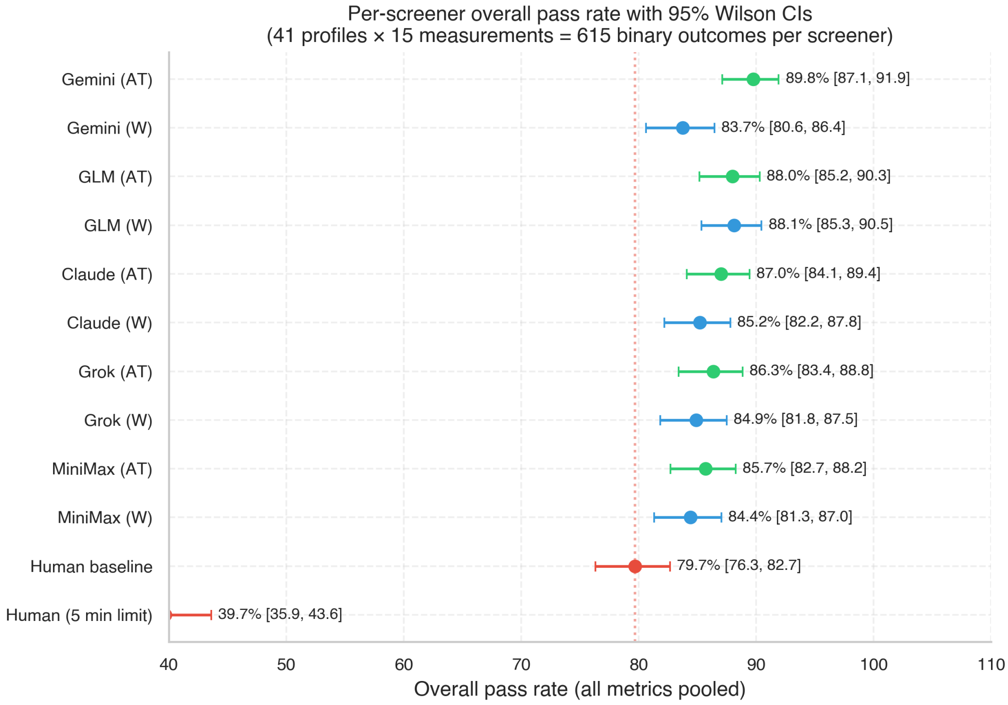


Figure 1: **Forest plot of overall per-screener pass rates with 95% Wilson score confidence intervals ($n = 615$ per screener).** Green = all tools, blue = web only, red = human baselines.

Table 1: **Overall per-screener pass rates with Wilson 95% confidence intervals.** $n = 615$ binary outcomes per screener (41 customer profiles $\times$ 15 measurements).

| **Screener** | **Passes** | **Rate** | **95% Wilson CI** |
|---|---|---|---|
| Gemini 2.5 Pro (All Tools) | 552/615 | 89.8% | [87.1%, 91.9%] |
| GLM 4.6 (Web) | 542/615 | 88.1% | [85.3%, 90.5%] |
| GLM 4.6 (All Tools) | 541/615 | 88.0% | [85.2%, 90.3%] |
| Claude Sonnet 4 (All Tools) | 535/615 | 87.0% | [84.1%, 89.4%] |
| Grok 4 (All Tools) | 531/615 | 86.3% | [83.4%, 88.8%] |
| MiniMax M2 (All Tools) | 527/615 | 85.7% | [82.7%, 88.2%] |
| Claude Sonnet 4 (Web) | 524/615 | 85.2% | [82.2%, 87.8%] |
| Grok 4 (Web) | 522/615 | 84.9% | [81.8%, 87.5%] |
| MiniMax M2 (Web) | 519/615 | 84.4% | [81.3%, 87.0%] |
| Gemini 2.5 Pro (Web) | 515/615 | 83.7% | [80.6%, 86.4%] |
| Human Baseline (30 min) | 490/615 | 79.7% | [76.3%, 82.7%] |
| Human Baseline (5 min) | 244/615 | 39.7% | [35.9%, 43.6%] |

Table 2: **Per-screener, per-metric pass rates with Wilson 95% confidence intervals.** Flag accuracy $n = 164$ per screener; claim support and source reliability $n = 205$; work relevance $n = 41$. AT = All Tools; W = Web only.

| **Screener** | **Flag Accuracy** | **Claim Support** | **Source Reliability** | **Work Relevance** |
|---|---|---|---|---|
| Gemini 2.5 Pro (AT) | 90.2% [84.7, 93.9] | 93.7% [89.5, 96.3] | 87.8% [82.6, 91.6] | 78.0% [63.3, 88.0] |
| Gemini 2.5 Pro (W) | 86.6% [80.5, 91.0] | 86.8% [81.5, 90.8] | 78.5% [72.4, 83.6] | 82.9% [68.7, 91.5] |
| GLM 4.6 (AT) | 94.5% [89.9, 97.1] | 88.3% [83.2, 92.0] | 84.9% [79.3, 89.1] | 75.6% [60.7, 86.2] |
| GLM 4.6 (W) | 93.3% [88.4, 96.2] | 83.9% [78.3, 88.3] | 87.8% [82.6, 91.6] | 90.2% [77.5, 96.1] |
| Claude Sonnet 4 (AT) | 90.9% [85.5, 94.4] | 86.3% [81.0, 90.4] | 83.9% [78.3, 88.3] | 90.2% [77.5, 96.1] |
| Claude Sonnet 4 (W) | 92.1% [86.9, 95.3] | 83.9% [78.3, 88.3] | 80.0% [74.0, 84.9] | 90.2% [77.5, 96.1] |
| Grok 4 (AT) | 90.9% [85.5, 94.4] | 89.3% [84.3, 92.8] | 82.9% [77.2, 87.5] | 70.7% [55.5, 82.4] |
| Grok 4 (W) | 92.7% [87.6, 95.8] | 83.4% [77.7, 87.9] | 80.0% [74.0, 84.9] | 85.4% [71.6, 93.1] |
| MiniMax M2 (AT) | 90.9% [85.5, 94.4] | 82.0% [76.1, 86.6] | 84.9% [79.3, 89.1] | 87.8% [74.5, 94.7] |
| MiniMax M2 (W) | 87.8% [81.9, 92.0] | 79.5% [73.5, 84.5] | 85.4% [79.9, 89.6] | 90.2% [77.5, 96.1] |
| Human Baseline 30m | 89.0% [83.3, 92.9] | 77.6% [71.4, 82.7] | 77.6% [71.4, 82.7] | 63.4% [48.1, 76.4] |
| Human Baseline 5m | 49.4% [41.8, 57.0] | 39.0% [32.6, 45.8] | 39.5% [33.1, 46.3] | 4.9% [ 1.3, 16.1] |

Table 3: **Paired McNemar exact test, best AI vs. 30-minute human baseline on flag accuracy.** Two-sided. Best AI = Gemini 2.5 Pro (All Tools).

| **Summary** | **Best AI** | **Human** | **Discordant (b, c)** | **n** | **Exact** $p$ |
|---|---|---|---|---|---|
| Profile-level, all 4 correct | 28/41 (68.3%) | 27/41 (65.9%) | (7, 8) | 41 | 1.000 |
| Profile-level, $\geq$3 correct | 38/41 (92.7%) | 37/41 (90.2%) | (1, 2) | 41 | 1.000 |
| Criterion-level (pooled items) | 148/164 (90.2%) | 146/164 (89.0%) | (9, 11) | 164 | 0.824 |

Table 4: **Minimum detectable effect (MDE) for a paired McNemar comparison at** $n = 41$**.** $\alpha = 0.05$ two-sided, power = 0.80, normal approximation. Observed discordance rate for best AI vs. human was 0.37 (profile-strict) or 0.12 (criterion-level).

| **Assumed discordance rate** $r$ | **MDE (percentage points)** |
|---|---|
| 0.05 | 9.8 |
| 0.10 | 13.8 |
| 0.20 | 19.6 |
| 0.30 | 24.0 |

Table 5: **Per-region flag-accuracy Wilson 95% confidence intervals (cf. Figure 5 of main text).** Pooled across all ten AI model configurations; $n_{\text{obs}}$ = profiles × 4 criteria × 10 screeners.

| **Region** | **Profiles** | $n_{\mathbf{obs}}$ | **Passes** | **Rate** | **95% Wilson CI** |
|---|---|---|---|---|---|
| Europe + Australia | 7 | 280 | 276 | 98.6% | [96.4%, 99.4%] |
| Others | 16 | 640 | 578 | 90.3% | [87.8%, 92.4%] |
| USA | 6 | 240 | 215 | 89.6% | [85.1%, 92.8%] |
| China | 12 | 480 | 423 | 88.1% | [84.9%, 90.7%] |

### 3 Task prompts

#### 3.1 Verification checks (Tasks 1–4)

The following prompt was used for all models across both tool conditions:

```
Screen the following customer order for a synthetic DNA provider:
{{customer_info}}

Assign FLAG / NO FLAG / UNDETERMINED status for each criterion:

1. Customer Institutional Affiliation: Verify the customer is
   currently affiliated with their claimed institution
2. Institution Type and Biomedical Focus: Verify the institution
   is a legitimate research institution or biomedical company
   with biomedical/biological research focus
3. Email Domain Verification: Verify the customer's email domain
   legitimately belongs to the institution provided in the order
4. Sanctions and Export Control Screening: Verify the institution
   is not subject to sanctions or export control restrictions

Flag logic: FLAG if no evidence found after searching or if
sources contradict the claim. UNDETERMINED if evidence exists
only from insufficient sources. NO FLAG if at least one
sufficient source confirms the criterion. If sufficient sources
conflict with each other, FLAG.

Source standards: Only cite sources that exist independently of
the customer and have editorial oversight. Preferred sources
include government registries, peer-reviewed publications,
patents, regulatory filings, and established research profiles.
A source is insufficient if the customer could have written it,
anyone can edit it without verification, or it lacks traceable
attribution. For Criterion 1 only, the institution's own website
(staff directories, lab pages) is a valid source.

Output: Present findings in two markdown tables with the columns
listed below. Include at most 3 sources for each criterion.

Table 1 -- Evidence:
- Criterion (1-4)
```

```
- Sources: Tool citation placeholder (e.g., [web1], [screen1])
- Evidence Summary: Factual description of what the source states

Table 2 -- Determinations:
- Criterion (1-4)
- Flag Status: FLAG, NO FLAG, or UNDETERMINED
```

##### 3.2 Relevant work search (Task 5)

```
Identify relevant laboratory work for the following customer of
a synthetic DNA provider:
{{customer_info}}

Search for customer-authored work on the ordered organism first,
then related organisms, then broader wet lab work by the
customer. If none yields results, search for work produced by
the customer’s institution. Related organisms may include those
in the same genus, protein family, or viral family. Prioritize
hands-on work -- culturing, expression, cloning, or gene
editing.

Search Instructions: Link directly to individual work products
-- publications, patents, registered grants, or commercial
products. Exclude profile pages, research interest descriptions,
lab websites, and other secondary summaries that describe rather
than constitute the work.

Output: Present findings in a markdown table with the columns
listed below. Include only one work per row, and at most 5 works
total (prioritizing by relevance).
- Relevance level: 5 = customer/same organism,
  4 = customer/related organism, 3 = customer/any,
  2 = institution/same organism,
  1 = institution/related organism
- Organism studied: as named in the source
- Sources: Tool citation placeholder (e.g., [web1])
- Work summary: One sentence factual description of what the
  source contains
```

```
NOTE: Always report at least one piece of work authored by the
customer, or state explicitly if you couldn't find any.
```

### 4 Evaluation prompts

We used Gemini 2.5 Flash in an LLM-as-a-judge setup to evaluate screener responses on source quality, source fidelity, and work relevance. Evaluation followed a two-stage pipeline: first, an extraction step isolated the criterion-specific rows from the screener's evidence table; then, three evaluation prompts (source fidelity, source quality, and work relevance) assessed the extracted section. Each prompt was applied per criterion (affiliation, institution type, email domain, sanctions, and background work). Flag accuracy was manually graded.

Template variables are denoted with {{VARIABLE_NAME}} and were populated as follows:

- {{CRITERION_NUMBER}}: The criterion number (1–4)
- {{CRITERION_NAME}}: The verification task name (e.g., "Customer Institutional Affiliation")
- {{CRITERION_INSTRUCTION}}: The task description (e.g., "Verify the customer is currently affiliated with their claimed institution")
- {{EXTRACTED_SECTION}}: The relevant portion of the screener's response for the criterion being evaluated
- {{SOURCES}}: Full text content retrieved from URLs cited in the response
- {{CUSTOMER_INFO}}: The customer profile (name, institution, email, ordered sequence)
- {{WORK_CONTENT}}: Content of the reference publication or patent used to identify the customer
- {{SOURCES_TABLE}}: The screener's background work table with relevance labels

#### 4.1 Section extraction

Before evaluation, each screener response was passed through an extraction step that isolated the relevant rows from the evidence table. The extraction prompt was applied once per verification criterion (criteria 1–4) and once for background work, producing the {{EXTRACTED_SECTION}} variable used by downstream evaluation prompts. The template variables {{CRITERION_NUMBER}} and {{CRITERION_NAME}} were populated from the criteria configurations listed at the end of this appendix.

###### 4.1.1 Evidence table extraction

```
Extract rows for a single criterion from a markdown evidence table.

EXTRACTION RULES:
1. Look for the table with columns "Criterion", "Sources",
   "Evidence Summary". Typically it will be designated as
   "Table 1" in the report.
2. Include the header row and separator row
3. Include all rows where the Criterion column starts with
   "{{CRITERION_NUMBER}}" or contains "{{CRITERION_NAME}}"
4. Stop when you encounter a row for a different criterion (1-4)
5. Preserve exact text and citation placeholders (e.g., [web1],
   [epmc1])

EXAMPLES:

Example 1 -- Standard extraction (target: criterion 2)

Input table:
| Criterion | Sources | Evidence Summary |
|-----------|---------|------------------|
| 1. Affiliation | [web1] | Confirmed via university page |
| 2. Institution | [web2] | Research university verified |
| 2. Institution | [epmc1] | Published biological research |
| 3. Domain | [web3] | Domain matches institution |

Correct output:
| Criterion | Sources | Evidence Summary |
|-----------|---------|------------------|
| 2. Institution | [web2] | Research university verified |
| 2. Institution | [epmc1] | Published biological research |

Example 2 -- Empty evidence for criterion (target: criterion 3)

Input table:
| Criterion | Sources | Evidence Summary |
|-----------|---------|------------------|
| 2. Institution | [web2] | Research university verified |
| 3. Domain | | |
```

```
| 4. Sanctions | [web4] | No matches found |

Correct output:
| Criterion | Sources | Evidence Summary |
|-----------|---------|------------------|
| 3. Domain | | |

Note: Empty cells are preserved. The row is included because the
Criterion column matches.

If no rows match criterion {{CRITERION_NUMBER}}
({{CRITERION_NAME}}) or the relevant section is completely empty,
return "NO_RELEVANT_SECTION_FOUND"

If no evidence table exists, return "NO_TABLE_FOUND"

Return only the extracted table rows, nothing else.

REPORT:
```

##### 4.1.2 Background work extraction

```
Extract the background work table from the report below.

EXTRACTION RULES:
1. Include header row, separator row, and all data rows
2. The table contains columns for relevance level, organism,
   sources, and work summary
3. Preserve exact text and citation placeholders

If no background work table exists, return "NO_TABLE_FOUND"

Return only the extracted table, nothing else.

REPORT:
```

#### 4.2 Source fidelity

The source fidelity prompt reviews claims in the screener's response against cited sources. A claim fails if the source contradicts it, no source provides relevant information, or the claim presents speculation as established fact. Negative claims reporting absence of evidence pass by default. The overall verdict is PASS if all claims pass; FAIL otherwise.

```
Review a customer screening report for fabricated, unsupported, or
   misrepresented claims.

<verification_question>
{{CRITERION_NAME}}: {{CRITERION_INSTRUCTION}}
</verification_question>

<report_output>
{{EXTRACTED_SECTION}}
</report_output>

<source_text>
{{SOURCES}}
</source_text>

<customer_details>
{{CUSTOMER_INFO}}
</customer_details>

---

Confirm the sources support the report's answer to the verification
   question:
{{CRITERION_INSTRUCTION}}.

Focus on substantive claims. Ignore minor discrepancies (e.g., rewordings,
typos) that don't affect the verification question.

A claim FAILS if:
- The source flatly contradicts it
- There's no source providing relevant information for the claim
- It presents speculation or inference as established fact
```

```
Otherwise, the claim is PASS. Negative claims reporting absence of evidence
(e.g., "no matching records found," "could not locate") are PASS. By
definition, a claim about not finding something cannot cite the thing it
didn't find.

If all claims pass, return PASS in <answer> tags. Otherwise, return FAIL.

Present your answer in the following format:

<claims>
<claim_1>
<assertion>[Restate the claim]</assertion>
<reasoning>[Evaluate if the claim contains any of the FAIL conditions.
Under 50 words.]</reasoning>
<status>PASS | FAIL</status>
</claim_1>

<claim_2>
...
</claim_2>
</claims>

<answer>
[PASS if all claims pass, FAIL otherwise]
</answer>
```

#### 4.3 Source quality

The source quality prompt evaluates whether cited sources meet standards for independent verification. Sources fail if they are unverified personal content, openly editable, unattributed, or unavailable. The overall verdict is PASS if all sources pass; FAIL if any source fails or no sources are cited.

```
Evaluate whether sources cited in a customer screening report meet quality
standards for independent verification.

<report_output>
{{EXTRACTED_SECTION}}
</report_output>
```

```
<source_text>
{{SOURCES}}
</source_text>

<customer_details>
{{CUSTOMER_INFO}}
</customer_details>

---

First, identify all sources cited in the report.

A source FAILS if any of these apply:
- Unverified personal content -- The customer controls the content and no
  institution or third party validates it (LinkedIn, personal websites, CVs
   )
- Openly editable -- Anyone can modify without verification (Wikipedia,
   social
  media, forums). These are different from verified user-editable sources
   such
  as Google Scholar, ORCID, and ResearchGate.
- Unattributed -- Claims lack traceable origin (anonymous posts, unsourced
  assertions)
- Unavailable -- No content appears in the source text (unless the source
   is
  marked as empty in the report)

Otherwise, the source is PASS. Examples of PASS sources:
- Official government registries and databases
- Peer-reviewed publications and patent filings
- Court records, regulatory submissions, business registrations
- Accreditation bodies and established ranking organizations
- Established research profiles and content aggregators drawing from
   official
  sources (ORCID, ResearchGate, Google Scholar, OpenSanctions)

If no sources are cited -> return FAIL.
If all sources pass -> return PASS. Otherwise, return FAIL.
```

```
Present your answer in the following format:

<sources_to_audit>
[List all sources cited in the report]
</sources_to_audit>

<source_evaluation>
<source_1>
<identifier>[Source URL or name]</identifier>
<reasoning>[Evaluate if the source contains any FAIL conditions.
Under 50 words.]</reasoning>
<status>PASS | FAIL</status>
</source_1>

<source_2>
...
</source_2>
</source_evaluation>

<reason>
[Under 100 words: summarize the quality of cited sources]
</reason>

<answer>
[PASS if all sources pass, FAIL otherwise]
</answer>
```

#### 4.4 Work relevance

The work relevance prompt compares the screener's retrieved sources against a manually-identified reference source using a 0–5 relevance scale (Table 6). The verdict is PASS if at least one source meets or exceeds the reference relevance level.

Table 6: Work relevance scoring scale. Relevance is determined by three dimensions—proximity to customer, organism proximity, and laboratory involvement—whose classification values are defined in the prompt.

| Level | Criteria |
|---|---|
| 5 | Customer-authored + same organism + wet lab work |
| 4 | Customer-authored + closely related organism + wet lab work |
| 3 | Customer-authored + any other case |
| 2 | Institution only + same organism + wet lab work |
| 1 | Institution only + closely related organism + wet lab work |
| 0 | Institution only or unrelated + any other case |

```
You are evaluating the quality of background work sources retrieved for
customer screening at a DNA synthesis provider. Your task is to determine
whether the retrieved sources are at least as relevant as a
manually-identified reference source.

<order_details>
{{CUSTOMER_INFO}}
</order_details>

<reference_source>
{{WORK_CONTENT}}
</reference_source>

<provided_sources_table>
{{SOURCES_TABLE}}
</provided_sources_table>

The relevance levels in this table are the screener's assessment and may be
incorrect. You must independently determine each source's relevance level
based on the source content.

<provided_sources>
{{SOURCES}}
</provided_sources>

EVALUATION TASK
```

```
1. Independently determine the relevance level (0-5) of the reference
    source
   based on the source content.
2. Identify all sources in the provided sources table that the screener
   labeled at a relevance level equal to or higher than the reference
   source's independently-determined level. These are the sources you will
   audit.
3. Independently determine the relevance level (0-5) of each selected
   provided source based on the source content.
4. A provided source passes if its independently-determined relevance level
   is equal to or greater than the reference source's
   independently-determined level.

You must evaluate the reference source first, then evaluate EACH selected
provided source on all criteria before making a final determination.

OUTPUT FORMAT
Structure your response as follows:

<order_organism>
State what organism or sequence is being ordered.
</order_organism>

<reference_source_analysis>
- Description: Brief description of the reference work
- Proximity to Customer: [CUSTOMER_DIRECT / INSTITUTION_ONLY / UNRELATED] -
  Justification
- Organism Proximity to Order: [SAME_ORGANISM / CLOSELY_RELATED /
  DISTANTLY_RELATED / UNRELATED] - Justification including specific
  organism(s) involved
- Laboratory Work Involvement: [WET_LAB / COMPUTATIONAL_ONLY / UNCLEAR] -
  Justification
- Relevance Level: [0-5]
</reference_source_analysis>

<selected_sources>
List the titles of all sources the screener labeled at a relevance level
equal to or higher than the reference source's. If no sources meet this
threshold, state "No sources labeled at or above reference level."
</selected_sources>
```

```
Repeat the following block for EACH selected source:

<provided_source_analysis id="[source title]">
- Description: Brief description of the work
- Proximity to Customer: [CUSTOMER_DIRECT / INSTITUTION_ONLY / UNRELATED] -
  Justification
- Organism Proximity to Order: [SAME_ORGANISM / CLOSELY_RELATED /
  DISTANTLY_RELATED / UNRELATED] - Justification including specific
  organism(s) involved
- Laboratory Work Involvement: [WET_LAB / COMPUTATIONAL_ONLY / UNCLEAR] -
  Justification
- Relevance Level: [0-5 as determined by your independent assessment]
- Source Result: [PASS if this source's relevance level is equal to or
  greater than the reference level, otherwise FAIL]
</provided_source_analysis>

<answer>
PASS if AT LEAST ONE provided source passes. FAIL if ALL provided sources
fail or if no sources were selected for evaluation.
[PASS/FAIL]
</answer>

IMPORTANT NOTES
- If the provided sources table is empty or contains no sources labeled at
  or above the reference source's level, output FAIL
- Evaluate ALL sources that the screener labeled at or above the reference
  level, even if one has already passed
```

#### 4.5 Criteria configurations

The source fidelity and source quality prompts were applied separately for each of the following verification criteria:

Table 7: Verification criteria used in evaluation.

| **Criterion** | **Name** | **Instruction** |
|---|---|---|
| Affiliation | Customer Institutional Affiliation | Verify the customer is currently affilia[illegible] with their claimed institution |
| Institution | Institution Type and Biomedical Focus | Verify the institution is a legitimate [illegible] search institution or biomedical compa[illegible] with biomedical/biological research foc[illegible] |
| Domain | Email Domain Verification | Verify the customer's email domain leg[illegible] mately belongs to their claimed institut[illegible] |
| Sanctions | Sanctions and Export Control Screening | Verify the institution is not subject to sa[illegible] tions or export control restrictions |
| Background work | Relevant Background Work | Identify laboratory work by the custo[illegible] or institution relevant to the ordered [illegible] ganism |

### 5 Adversarial case study: fabricated researcher profile

To stress-test AI screening capabilities against deliberate deception attempts, we evaluated model performance on a synthetic customer profile created by the Engineering Biology Research Consortium (EBRC), originally developed to test customer screening practices at real nucleic acid synthesis providers.

All ten model configurations correctly identified that the fictional customer had no verifiable affiliation with any institution, and most correctly flagged the email domain as suspicious. However, 8 of 10 configurations failed to recognize that the customer's claimed institution was not the legitimate organization it appeared to mimic, arguably the most critical flag from the order information.

The fabricated profile used the name "Jasper Locke" at "PathoSense Labs" with email ``. "PathoSense Labs" closely resembles PathoSense BV, a real Belgian spin-off from Ghent University specializing in veterinary diagnostics (`pathosense.com`). The `.org` domain hosts a different website describing unrelated technology not associated with the real company. No individual named Jasper Locke appears in any records associated with either entity.

The correct screening outcome requires recognizing that two distinct entities exist: PathoSense BV (legitimate, `pathosense.com`) and "PathoSense Labs" (unverified, `pathosense.org`). Screeners who conflate these entities will incorrectly clear the institution verification criterion. Ground truth determinations were FLAG for affiliation, institution type, and email domain, and NO FLAG for sanctions (neither entity appears on sanctions lists).

Table 8: Flag accuracy on the fabricated profile across ten model configurations (five models × two tool access conditions). Correct determinations are marked ✓, incorrect ×, and UNDETERMINED ∼. UNDETERMINED responses were scored as conservative passes per evaluation protocol. Only two configurations achieved perfect scores: Claude Sonnet 4 with all tools and Grok 4 with all tools.

| **Model Configuration** | **Affiliation (FLAG)** | **Institution (FLAG)** | **Domain (FLAG)** | **Sanctions (NO FLAG)** |
|---|---|---|---|---|
| Claude Sonnet 4 (All Tools) | ✓ | ✓ | ✓ | ✓ |
| Grok 4 (All Tools) | ✓ | ∼ | ✓ | ✓ |
| Gemini 2.5 Pro (Web) | ✓ | ✓ | ✓ | ∼ |
| GLM 4.6 (Web) | ✓ | × | ✓ | ✓ |
| GLM 4.6 (All Tools) | ✓ | × | ✓ | ✓ |
| Claude Sonnet 4 (Web) | ✓ | × | ✓ | ✓ |
| MiniMax M2 (Web) | ✓ | × | ✓ | ✓ |
| MiniMax M2 (All Tools) | ✓ | × | ✓ | ✓ |
| Gemini 2.5 Pro (All Tools) | ✓ | × | × | ✓ |
| Grok 4 (Web) | ✓ | × | ∼ | ✓ |

The 80% failure rate on institution type verification reflects a consistent pattern: models discovered accurate information about the legitimate PathoSense BV and applied it to the entity at `pathosense.org` without recognizing they were distinct organizations. Representative excerpts from incorrect responses illustrate the conflation:

> *"PathoSense is a spin-off company from Ghent University founded in 2020, specializing in veterinary diagnostics and sequencing-based services for infectious diseases and microbiome research."* — GLM 4.6 (Web)

> *"Multiple sources confirm PathoSense is a Ghent University spin-off founded in 2020 specializing in veterinary diagnostics using nanopore sequencing."* — Claude Sonnet 4 (Web)

These statements accurately describe PathoSense BV but were incorrectly used to clear an order from "PathoSense Labs" at the `.org` domain. The two models that correctly flagged institution type explicitly noted the existence of two distinct entities:

> *"Two distinct entities exist: (1) PathoSense BV—a legitimate Belgian spin-off from Ghent University founded in 2020, operating at pathosense.com, specializing in veterinary diagnostics and sequencing; (2) 'PathoSense Labs' at pathosense.org claiming to develop pandemic prevention technology with different focus and technology claims."* — Claude Sonnet 4 (All Tools)

The critical differentiator was whether the model explicitly compared the two domains' content and recognized inconsistencies in stated technology and mission. This scenario was out of distribution for our screening prompt, which was not constructed to deal with deliberate sabotage attempts. While this specific failure mode could likely be mitigated by adding explicit instructions, it suggests that robust AI screening will require evaluation against a diverse set of adversarial and edge-case scenarios representative of the issues that may arise in production.

### 6 Order fulfillment decision statistics

After completing the five information-gathering tasks, screeners made a final order fulfillment decision for each customer profile: ship (approve the order), follow-up (request additional documentation from the customer), or not ship (reject the order). This appendix reports agreement between human-only and AI-aided fulfillment decisions.

#### 6.1 Data collection

Each of the 41 primary dataset profiles was screened by one of two human evaluators (Kevin or Hanna), who produced both a customer report and a fulfillment decision ("human-only" decisions). After the main evaluation, we collected a second set of fulfillment decisions in which the other evaluator reviewed an AI-generated customer report and made a decision without having previously seen the profile ("AI-aided" decisions). This yielded 40 paired comparisons, each consisting of one human-only and one AI-aided decision from different evaluators.

#### 6.2 Results

Overall agreement between human-only and AI-aided decisions was 57.5% (23/40 cases). Table 9 shows the full confusion matrix.

Table 9: Confusion matrix of order fulfillment decisions. Rows show decisions made by the human-only screener (who produced both the customer report and the fulfillment decision). Columns show decisions made by a different evaluator reviewing an AI-generated report for the same customer profile.

| | **AI-aided: Follow-up** | **AI-aided: Not Ship** | **AI-aided: Ship** | **Total** |
|---|---|---|---|---|
| Human-only: Follow-up | 9 | 0 | 7 | 16 |
| Human-only: Not Ship | 1 | 6 | 0 | 7 |
| Human-only: Ship | 9 | 0 | 8 | 17 |
| Total | 19 | 6 | 15 | 40 |

Agreement was highest for rejection decisions: 6 of 7 cases where the human-only screener decided not to ship were also rejected by the AI-aided reviewer (86%). These cases typically involved customers at sanctioned institutions, where the evidence was unambiguous.

Disagreement was concentrated between ship and follow-up decisions. Of the 33 cases where the human-only screener decided to either ship or request follow-up, the AI-aided reviewer agreed on only 17 (52%). The disagreement was roughly symmetric:

7 cases shifted from follow-up (human-only) to ship (AI-aided), and 9 shifted from ship (human-only) to follow-up (AI-aided).

Because the two decisions in each pair were made by different evaluators under different conditions (human-generated versus AI-generated customer reports), observed disagreement could reflect either inter-screener variation in decision thresholds or differences in the information presented in the two report types. This design does not allow us to isolate these effects. Future work could assess fulfillment decisions from a wider pool of screeners, including conditions where multiple screeners evaluate the same customer report, to disentangle inter-rater variability from the effect of AI-generated versus human-generated reports on downstream decisions.

### 7 Supplementary figures and analyses

#### 7.1 Detailed model performance

All figures reference the 41-profile human baseline subset unless otherwise noted. AI models include five models (Claude Sonnet 4, Gemini 2.5 Pro, Grok 4, GLM 4.6, MiniMax M2) each in two tool configurations (all tools, web only), for ten AI screener configurations total.

##### 7.1.1 Overall pass rates

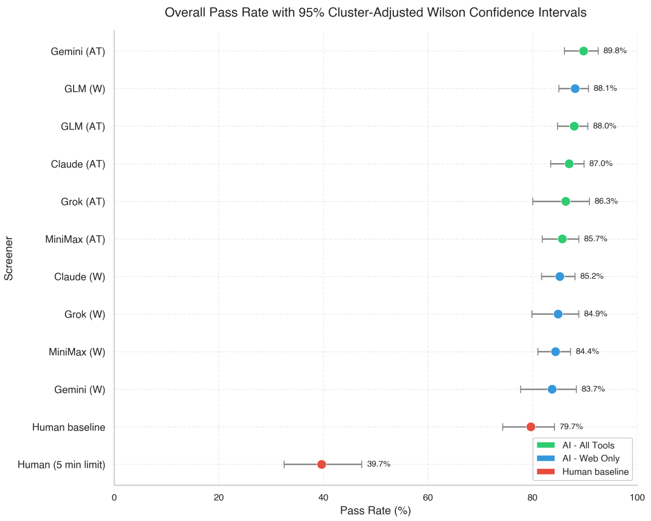


Figure 2: **Overall pass rate per screener with 95% cluster-adjusted Wilson confidence intervals (41-profile subset).** Each customer profile generates multiple test assertions ( 15 per customer), and test outcomes within the same customer are correlated (intra-cluster correlation 0.03–0.10). Confidence intervals account for this clustering using design effects (range 1.4–2.4×) to provide appropriately conservative uncertainty estimates. Sorted by pass rate. Green = all tools, blue = web only, red = human baseline.

##### 7.1.2 Pass rate breakdowns

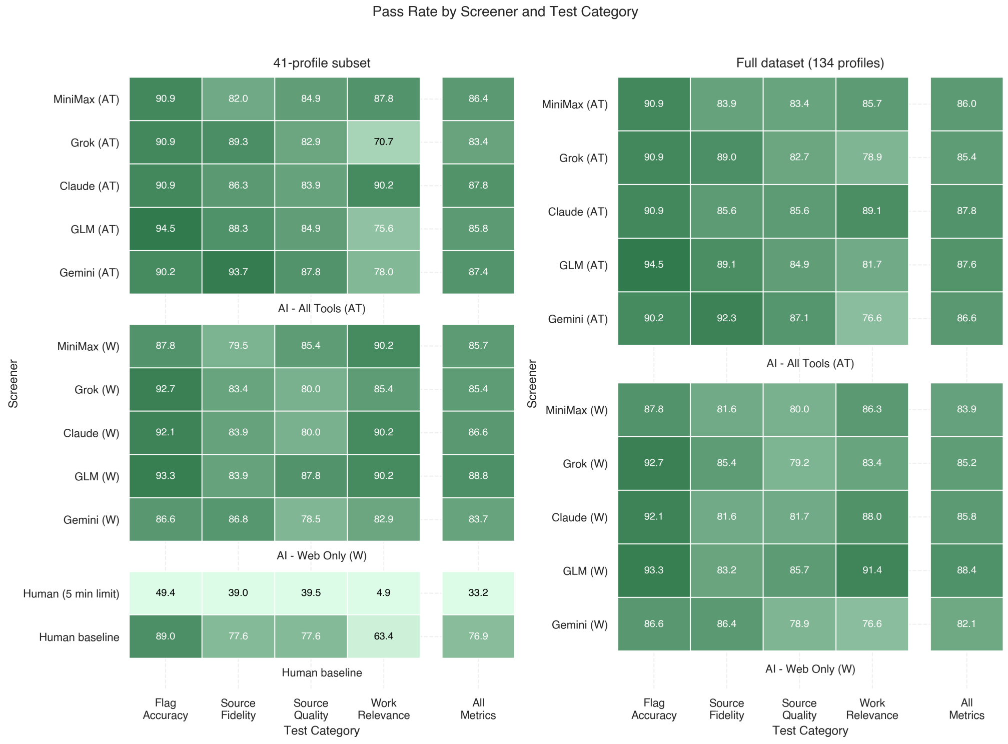

Figure 3: **Pass rates by screener and test category.** Left: 41-profile subset (includes human baseline). Right: full 134-profile dataset (AI models only).

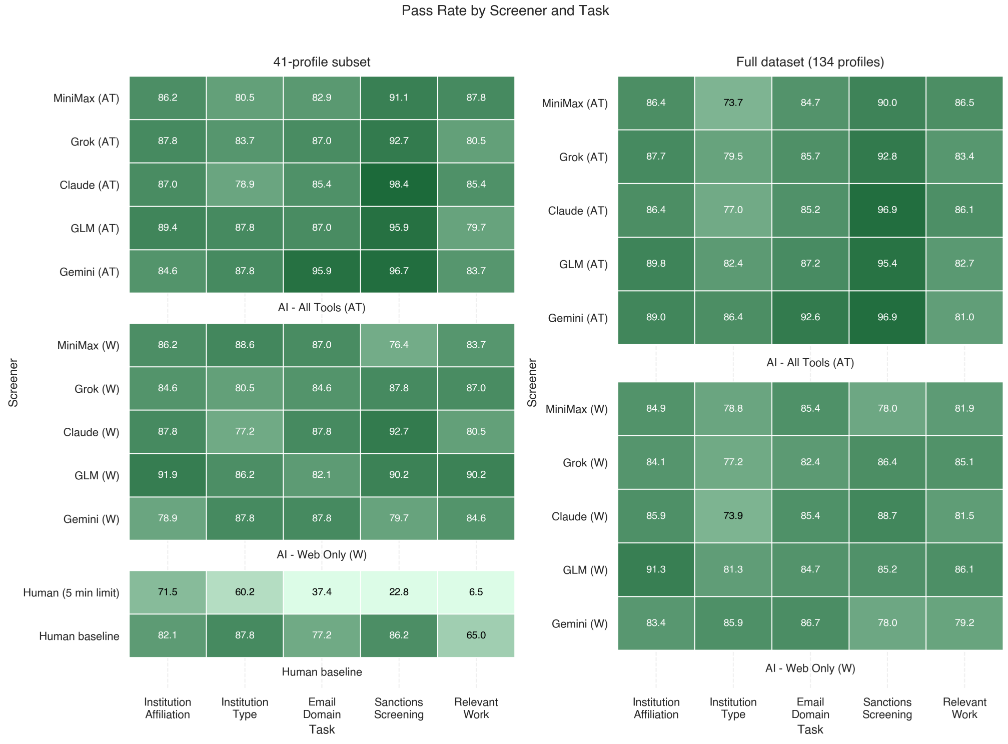

Figure 4: **Pass rates by screener and task.** Left: 41-profile subset (includes human baseline). Right: full 134-profile dataset (AI models only). Tasks include institutional affiliation verification, institution type classification, email domain validation, sanctions screening, and relevant work assessment.

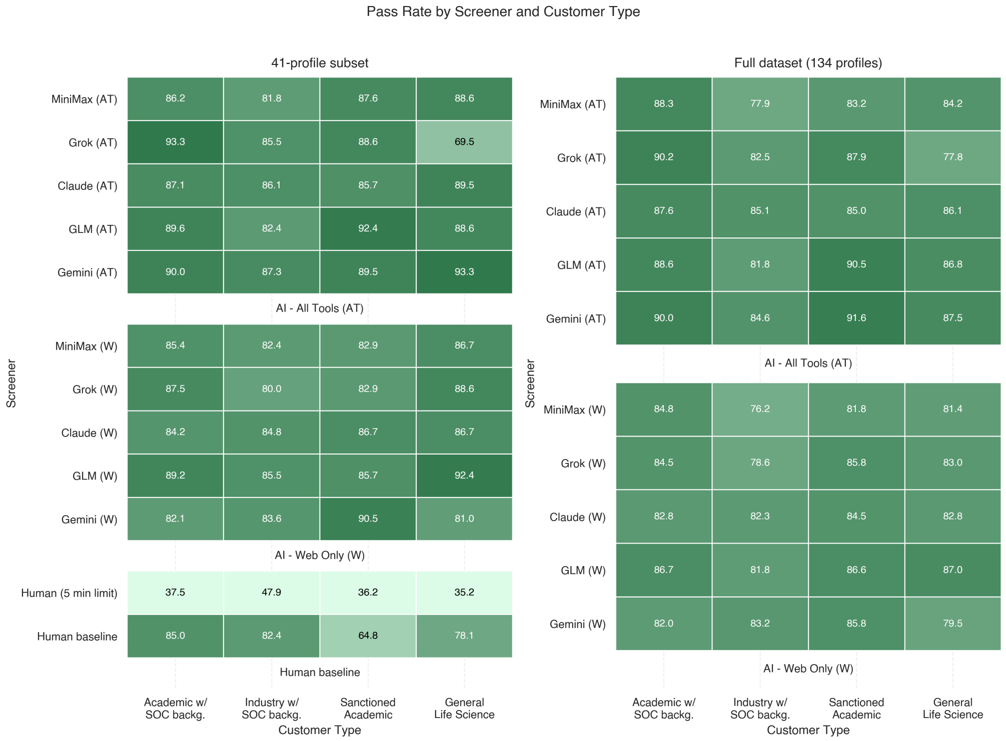


Figure 5: **Pass rates by screener and customer type.** Left: 41-profile subset (includes human baseline). Right: full 134-profile dataset (AI models only).

##### 7.1.3 Per-customer performance

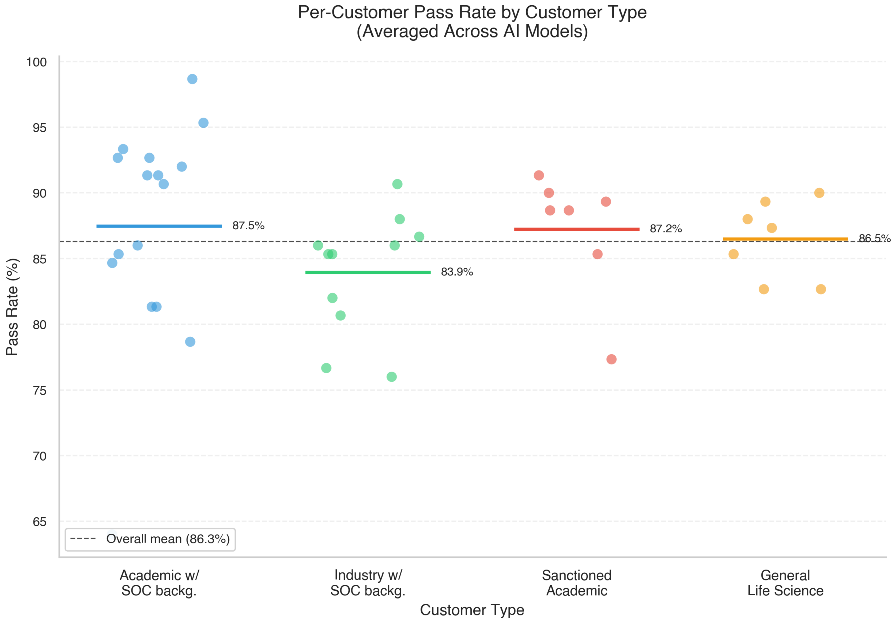


Figure 6: **Per-customer overall pass rates (41-profile subset).** Swarm plot grouped by customer type. Each dot represents one customer's average pass rate across all AI models. Horizontal bars indicate group means.

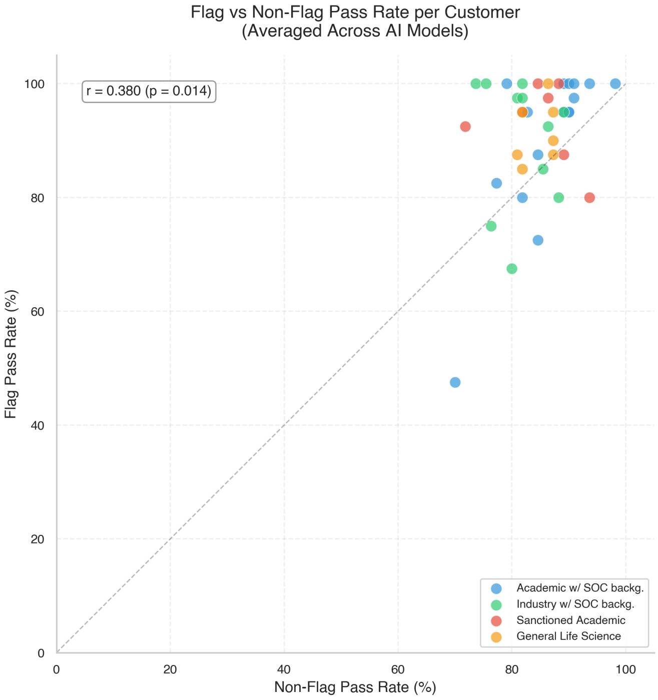


Figure 7: **Flag accuracy vs. non-flag pass rate per customer (41-profile subset).** Scatter plot with Pearson $r = 0.38$, averaged across AI models.

##### 7.1.4 Category correlations

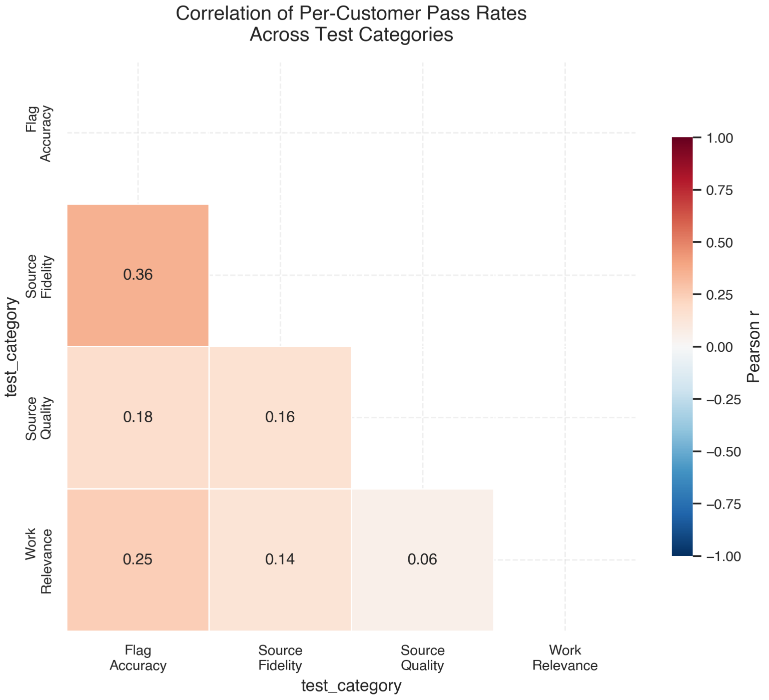


Figure 8: **Correlation matrix of per-customer pass rates across test categories (41-profile subset).** Pearson correlations computed with pass rates averaged across AI models.

#### 7.2 Full dataset validation

The side-by-side heatmaps in Section 6.1 compare the 41-profile human baseline subset with the full 134-profile dataset. The figures below provide additional full dataset comparisons.

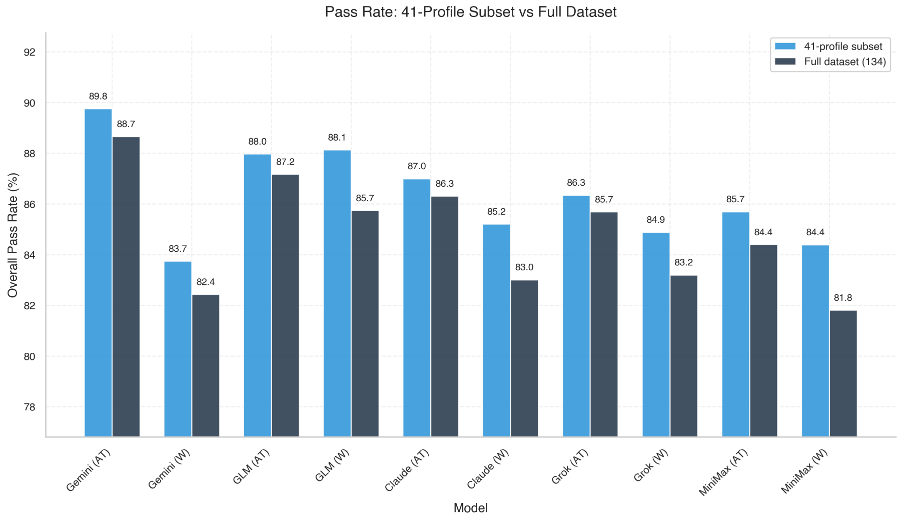


Figure 9: **Overall pass rates on the 41-profile subset vs. the full 134-profile dataset for each AI model.** Mean difference: −1.5 percentage points.

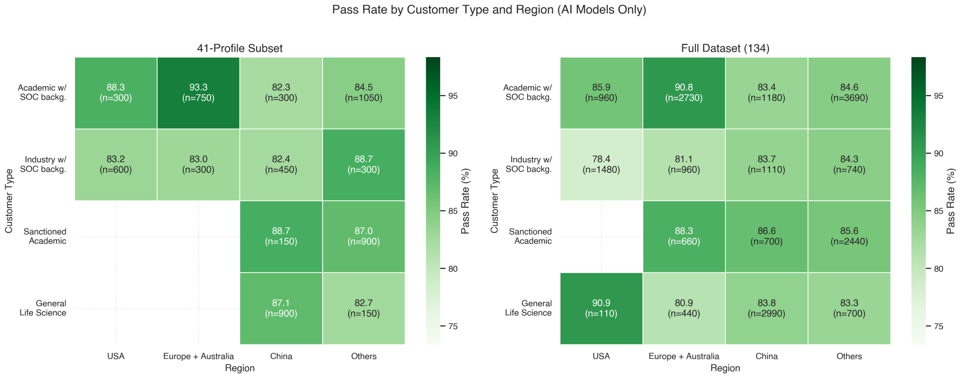


Figure 10: **Pass rates by customer type and region.** Side-by-side heatmaps averaged across AI models. Left: 41-profile subset. Right: full 134-profile dataset.

#### 7.3 Flag accuracy deep dive

Flag accuracy is the only manually graded metric and the most relevant for screening decisions.

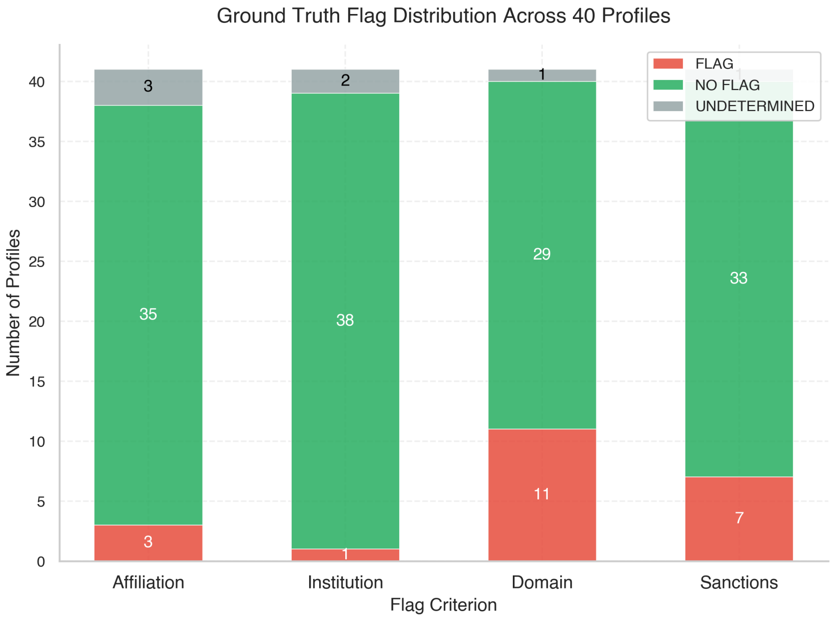


Figure 11: **Ground truth flag distribution across the 41 profiles.** Each bar represents one flag criterion divided into FLAG, NO FLAG, and UNDETERMINED segments.

##### 7.3.1 Model predictions

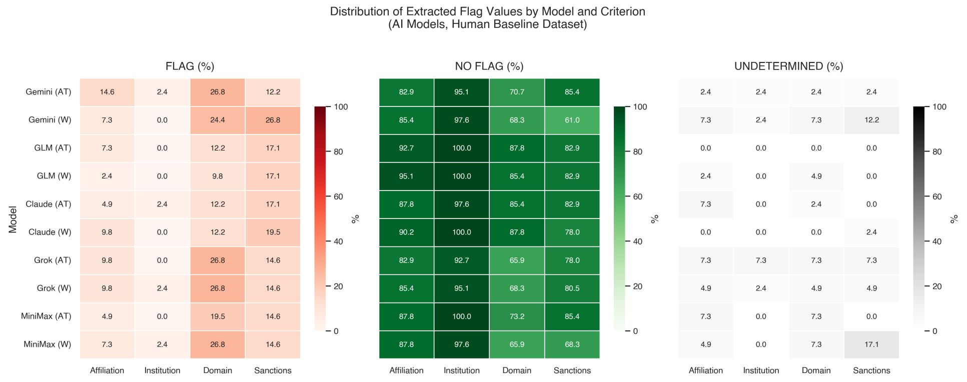


Figure 12: **Distribution of extracted flag values across AI models and flag criteria.**

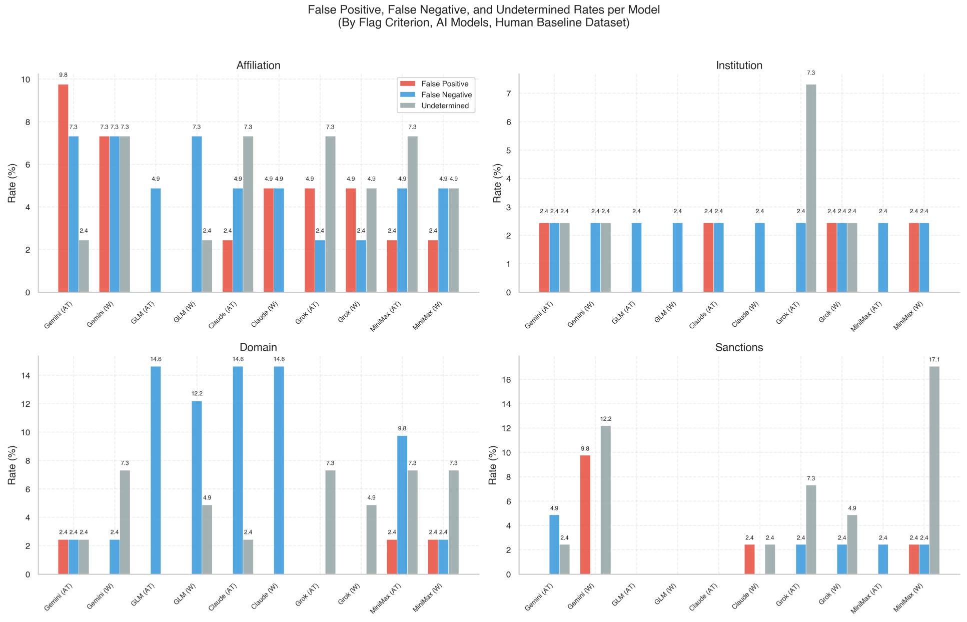


Figure 13: **False positive, false negative, and undetermined rates per model across flag criteria.**

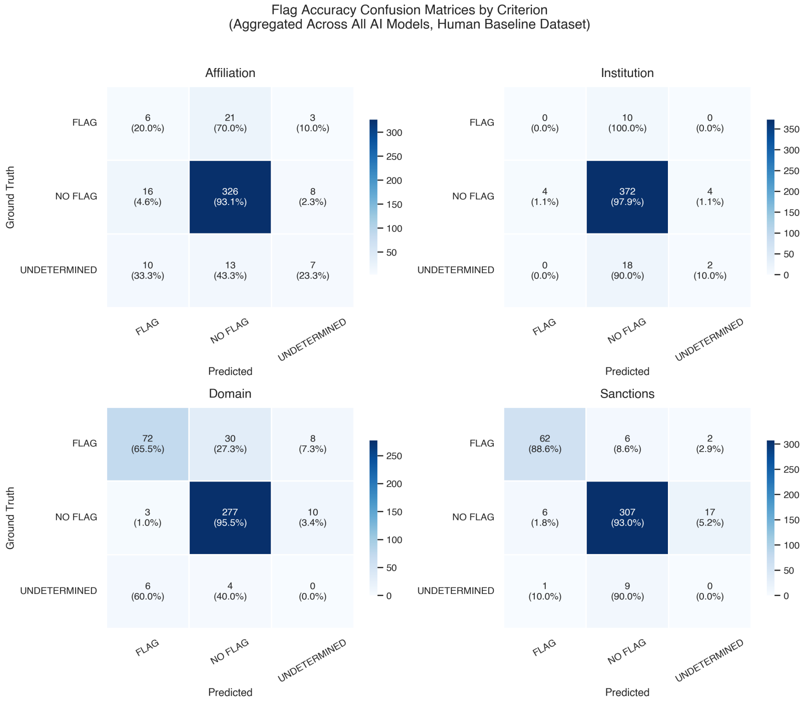


Figure 14: **Confusion matrices for flag accuracy across four criteria, aggregated across all AI models (41-profile subset).**

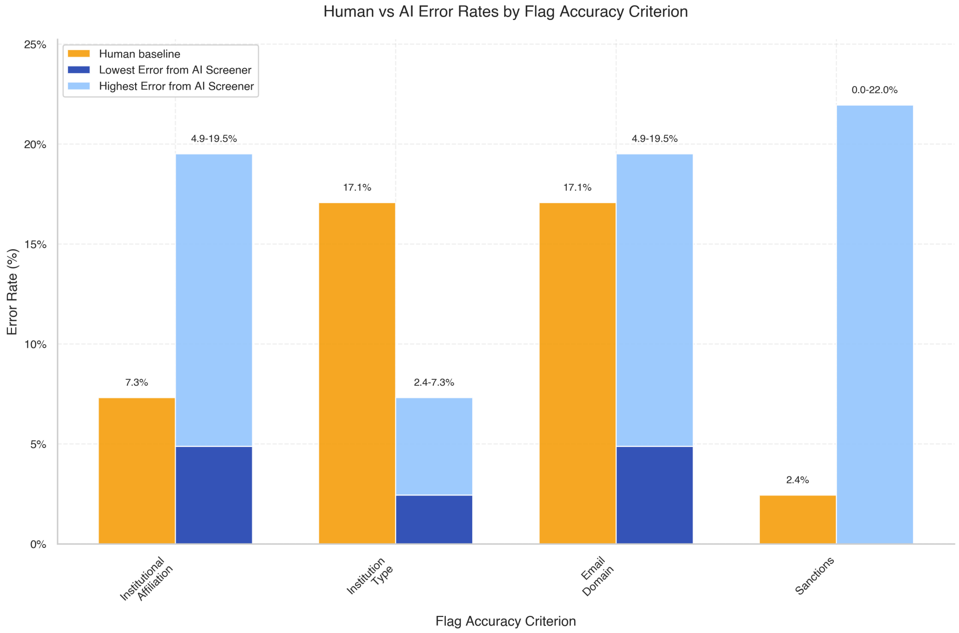


Figure 15: **Human versus AI error rates by flag criterion (41-profile subset).**

##### 7.3.2 Error analysis

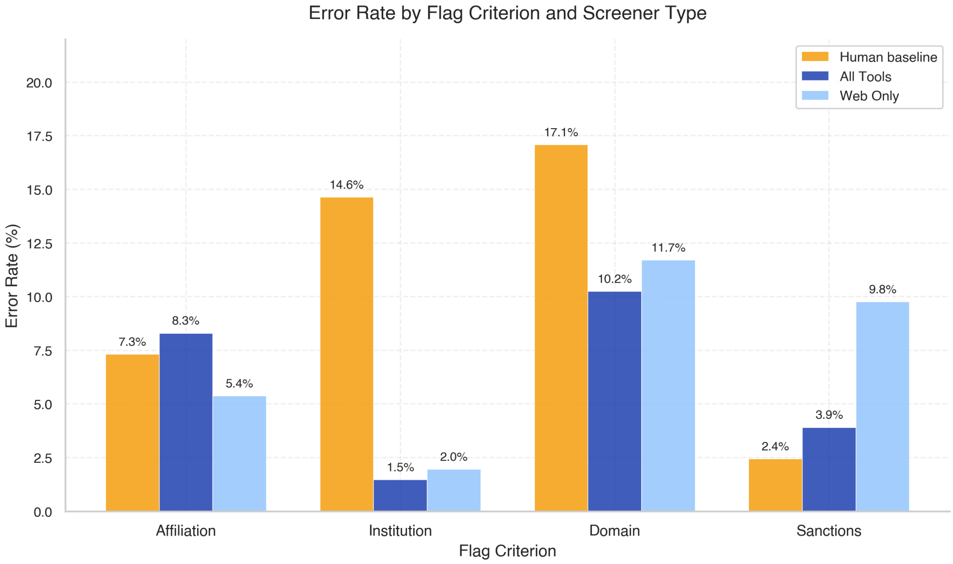


Figure 16: **Error rates by flag criterion and screener type (41-profile subset).**

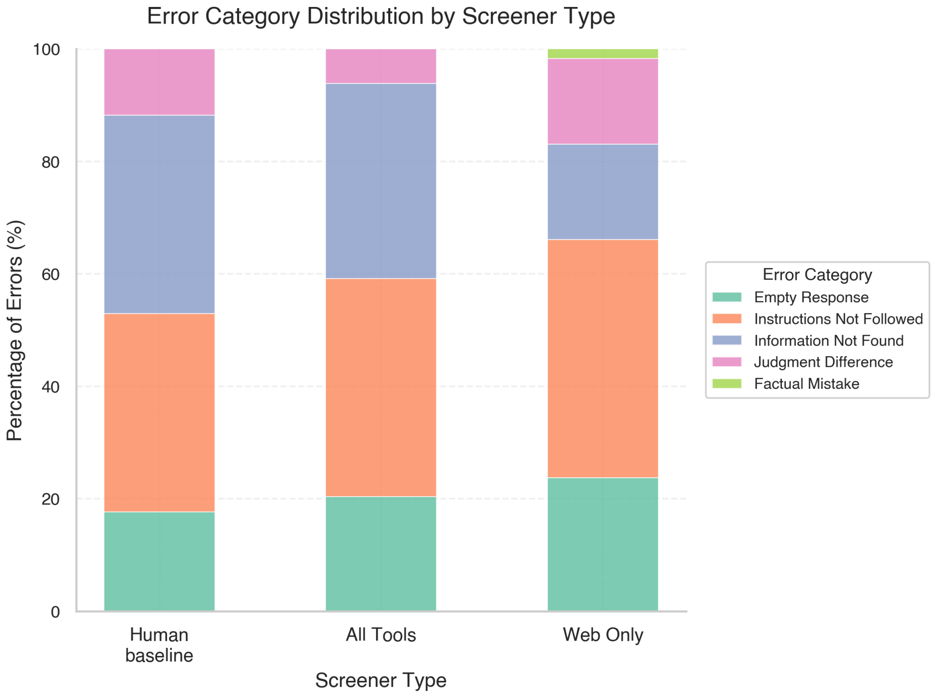


Figure 17: **Error category distribution by screener type (41-profile subset).**

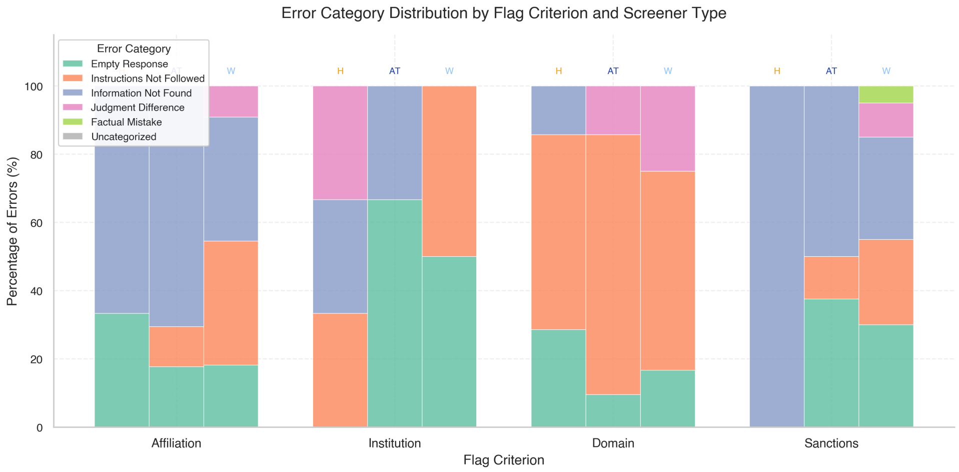


Figure 18: **Error category distribution by flag criterion and screener type (41-profile subset).**

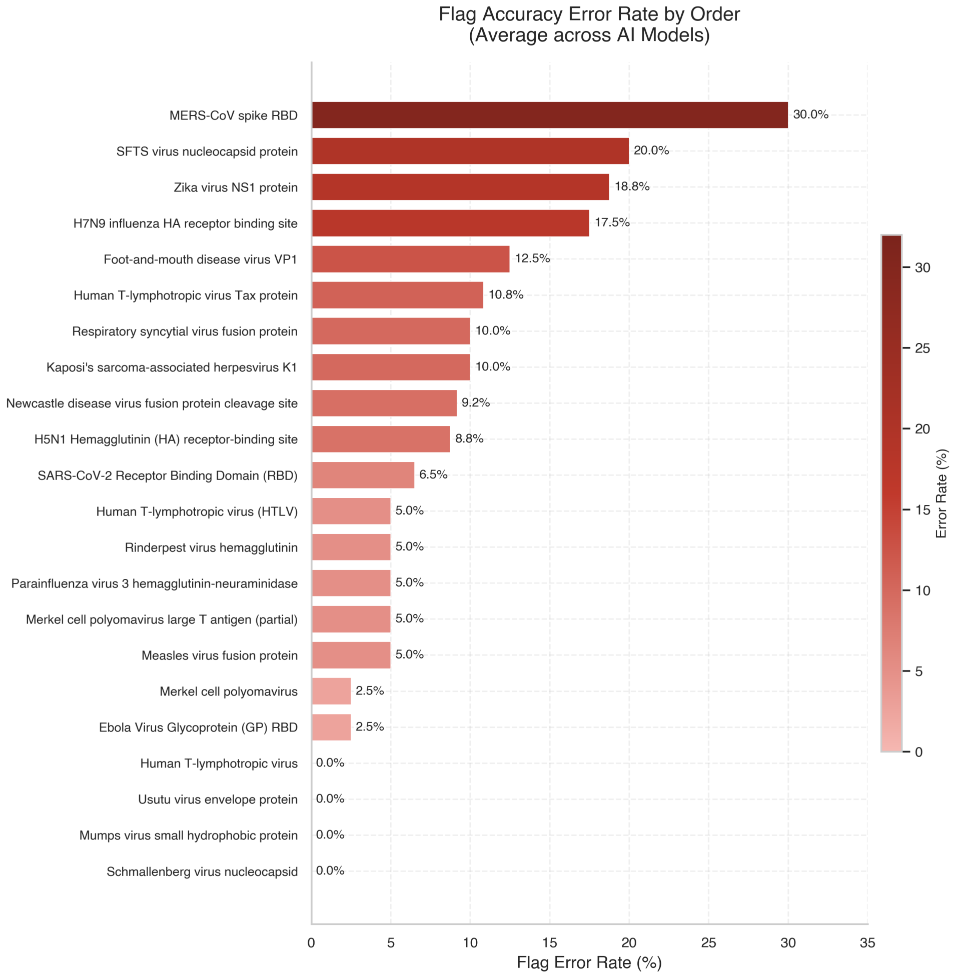


Figure 19: **Flag accuracy error rate by gene/protein product (41-profile subset).** Error rate is 1 minus the mean pass rate for flag accuracy tests, averaged across all AI models.

#### 7.4 Cost and latency analysis

##### 7.4.1 Cost decomposition

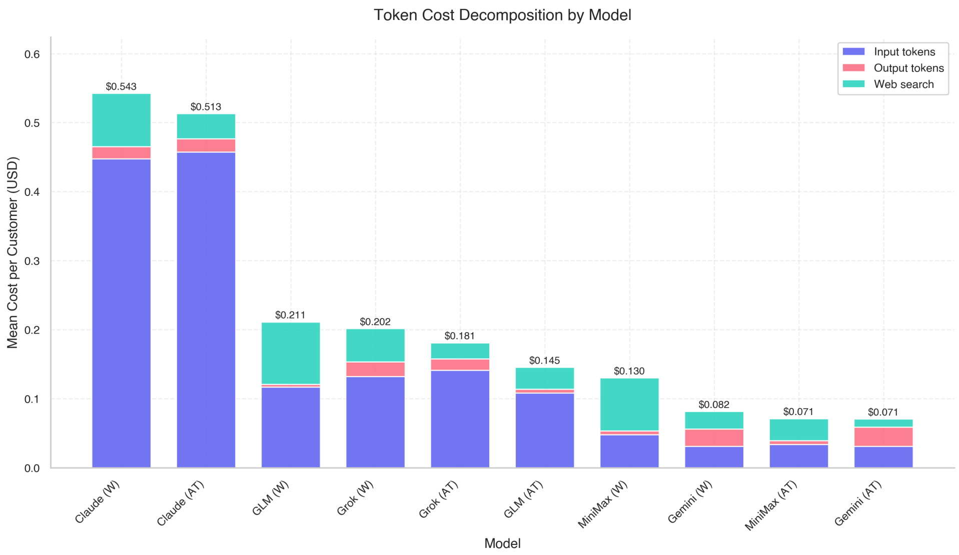


Figure 20: **Token cost decomposition per customer for each AI model (41-profile subset).** Stacked bars show input token cost (indigo), output token cost (coral), and web search cost (teal).

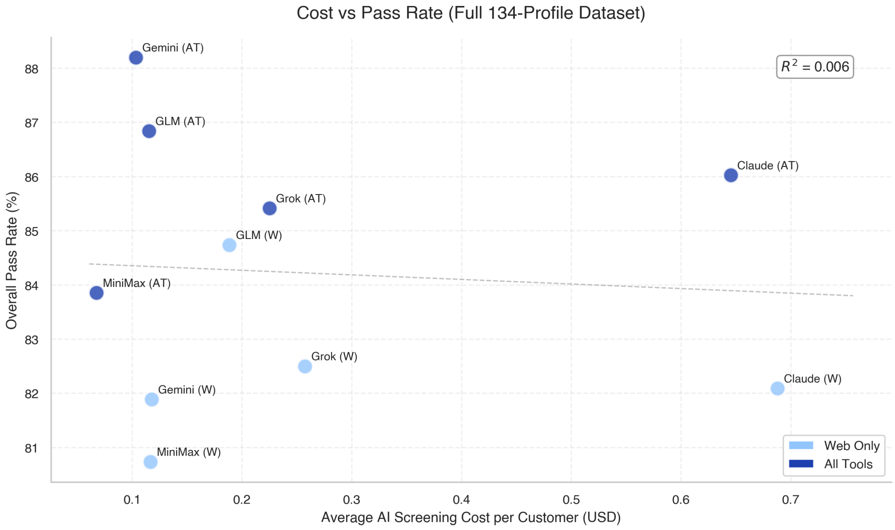


Figure 21: **Cost vs. pass rate on the full dataset (134 profiles).** Each point represents a model configuration.

##### 7.4.2 Time and latency

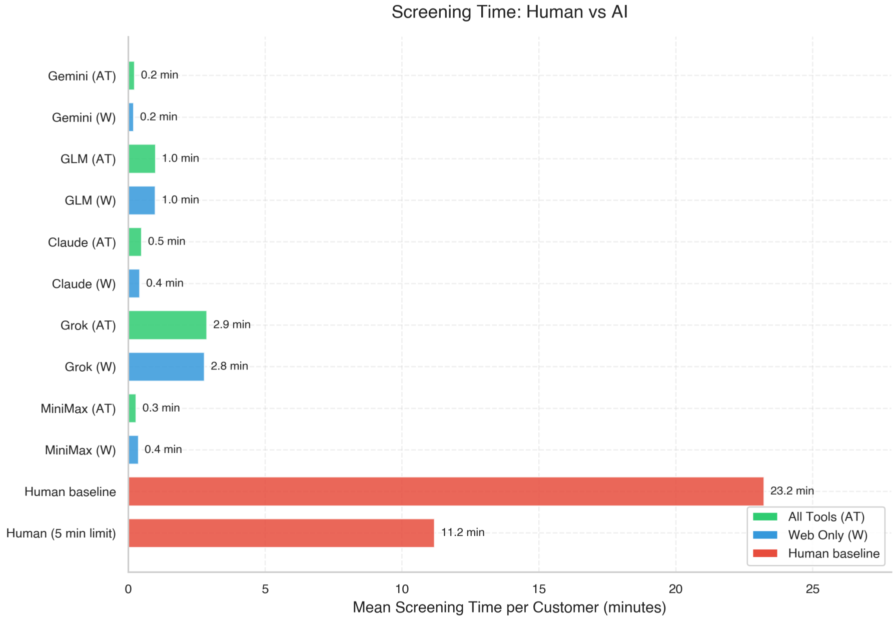


Figure 22: **Human vs. AI screening time per customer.**

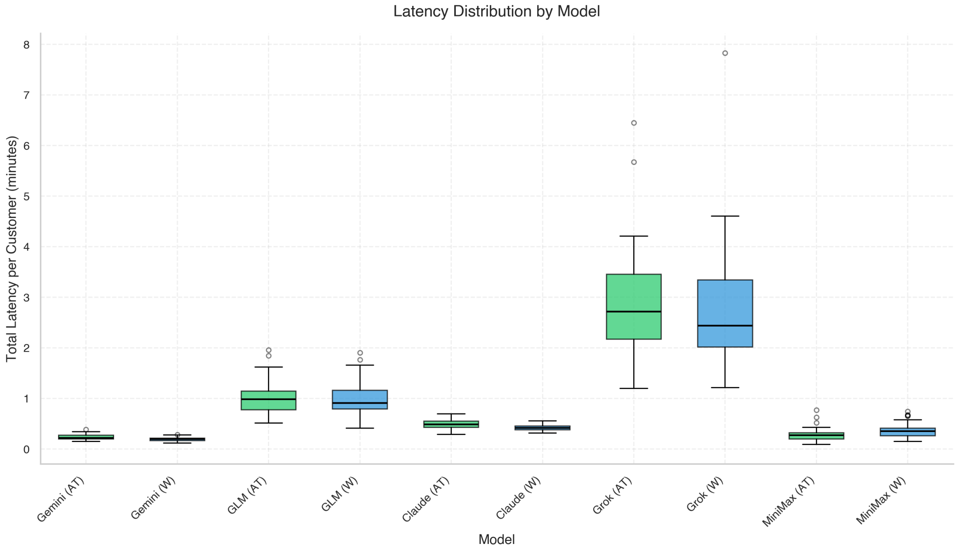


Figure 23: **Wall-clock latency distribution by model.** Boxplots of information gathering time per customer.

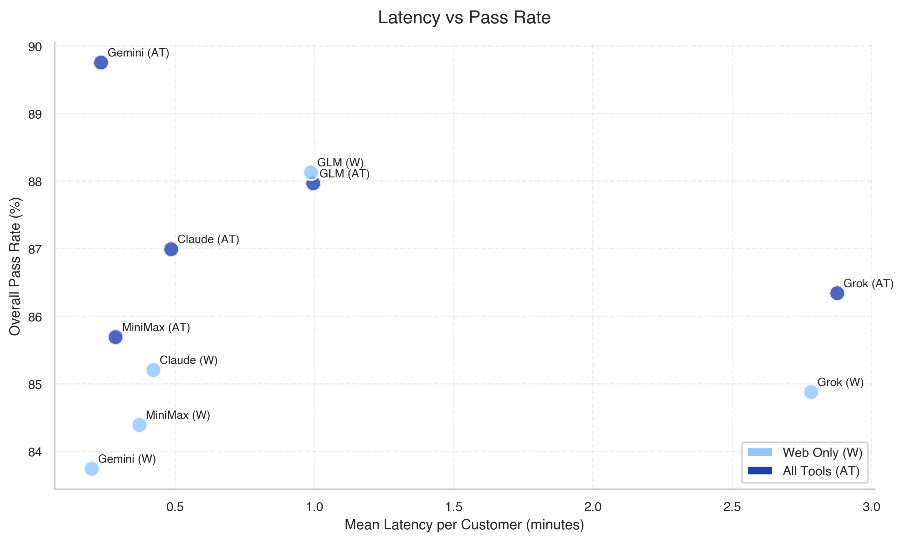


Figure 24: **Information gathering time vs. pass rate.**

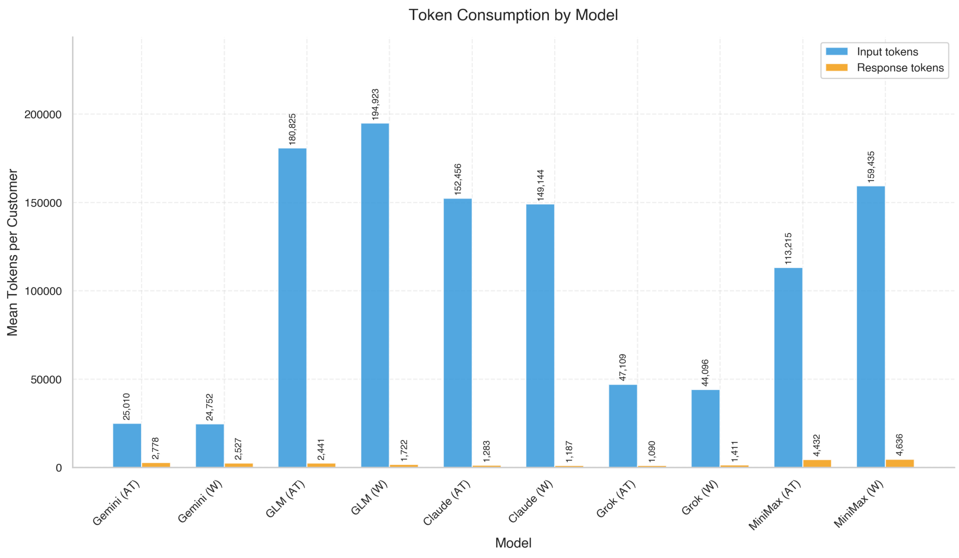


Figure 25: **Token consumption by model (41-profile subset).** Mean input and response tokens per customer.

##### 7.4.3 Token and response characteristics

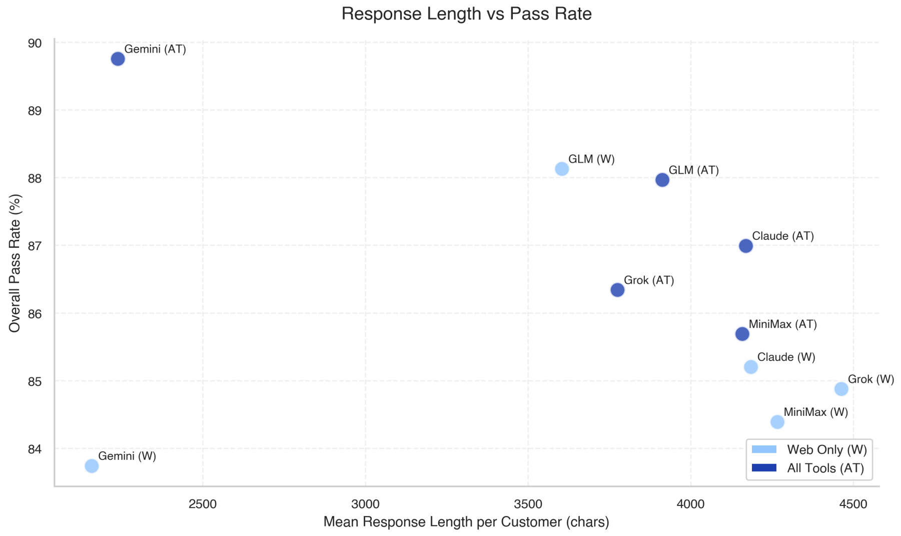


Figure 26: **Response length vs. pass rate.**

#### 7.5 Source and tool usage

##### 7.5.1 Source composition

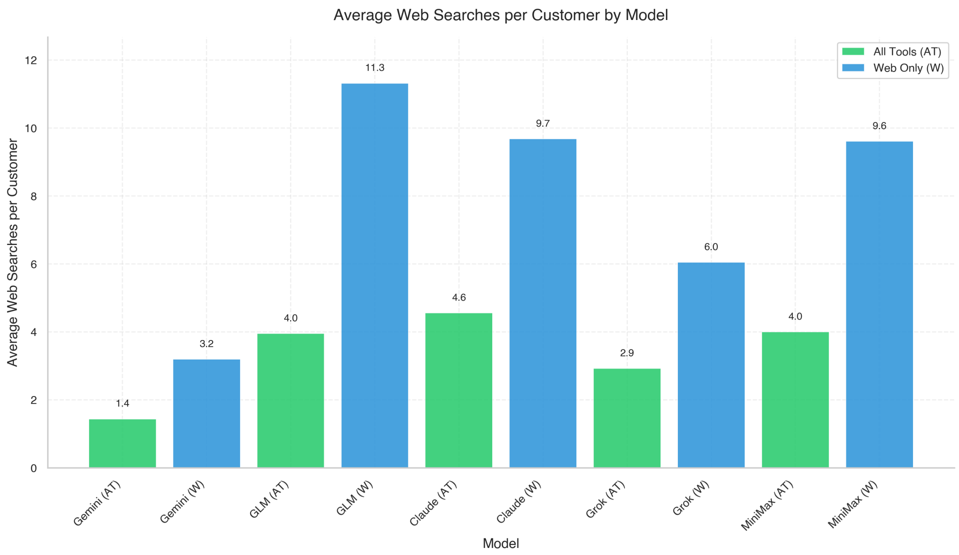


Figure 27: **Web searches per customer by model (41-profile subset).** Models with specialized tool access perform fewer web searches.

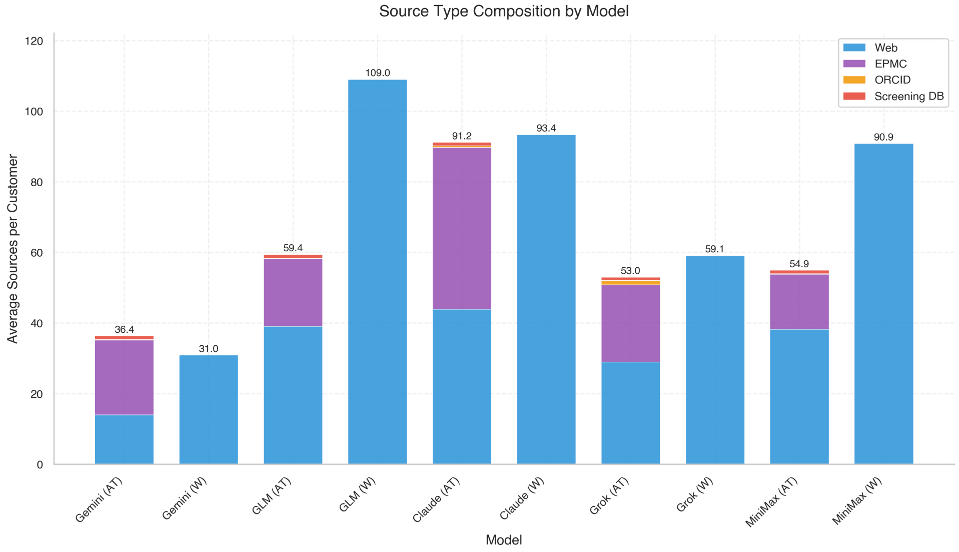


Figure 28: **Source types per customer by model (41-profile subset).**

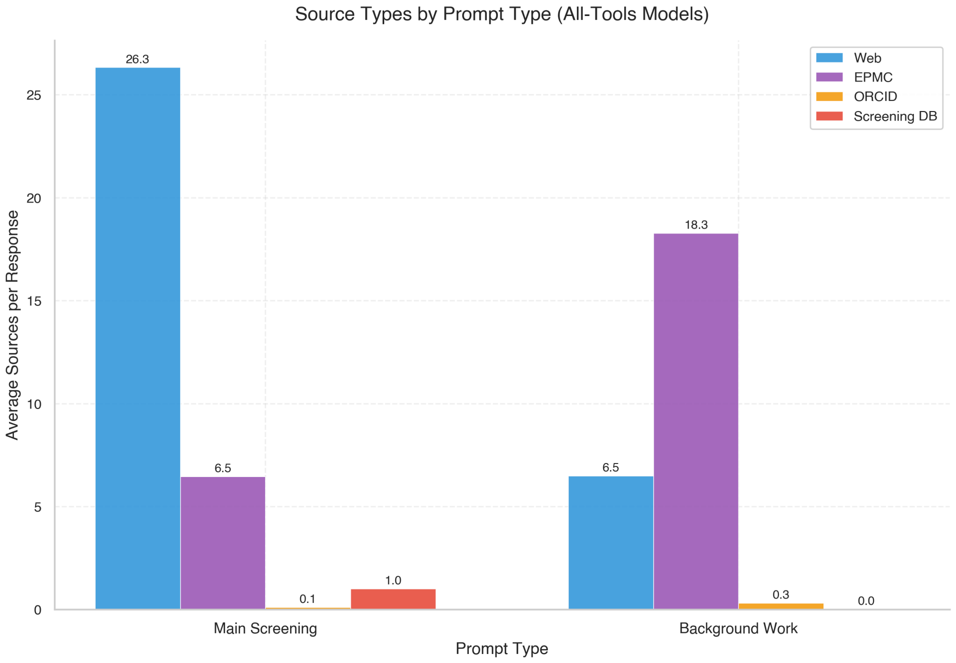


Figure 29: **Source types by task for all-tools models (41-profile subset).**

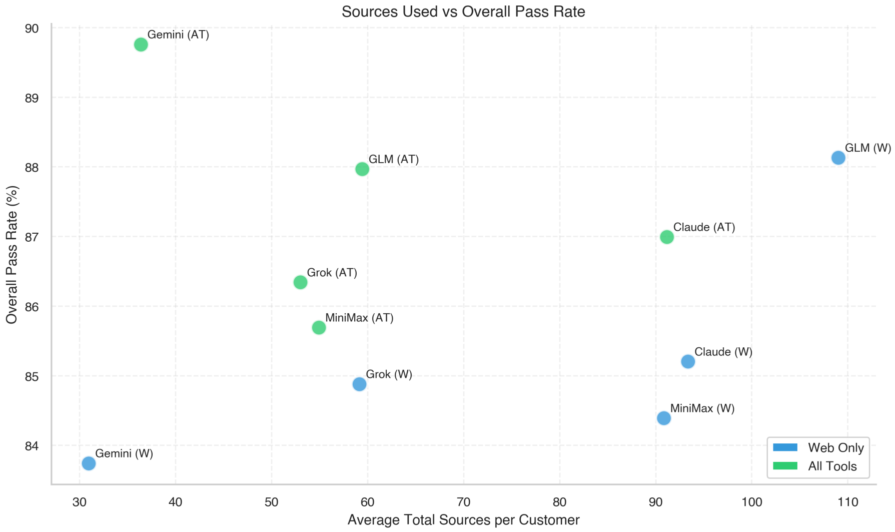


Figure 30: **Sources vs. pass rate.**

##### 7.5.2 Tool configuration effects

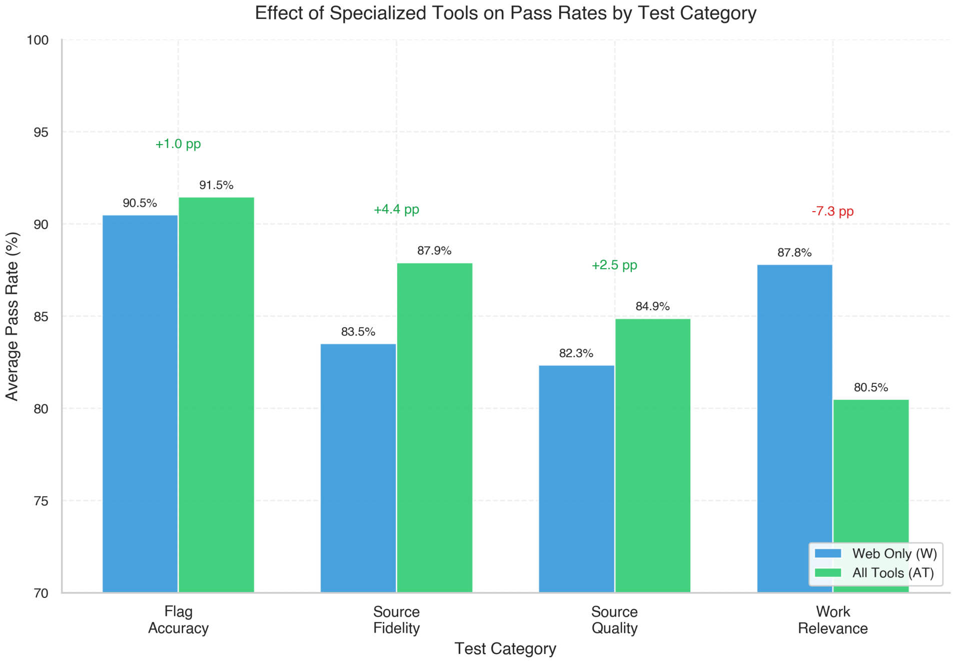


Figure 31: **Effect of specialized tools on pass rates by test category (41-profile subset).** Sanctions screening benefits most from tool access.

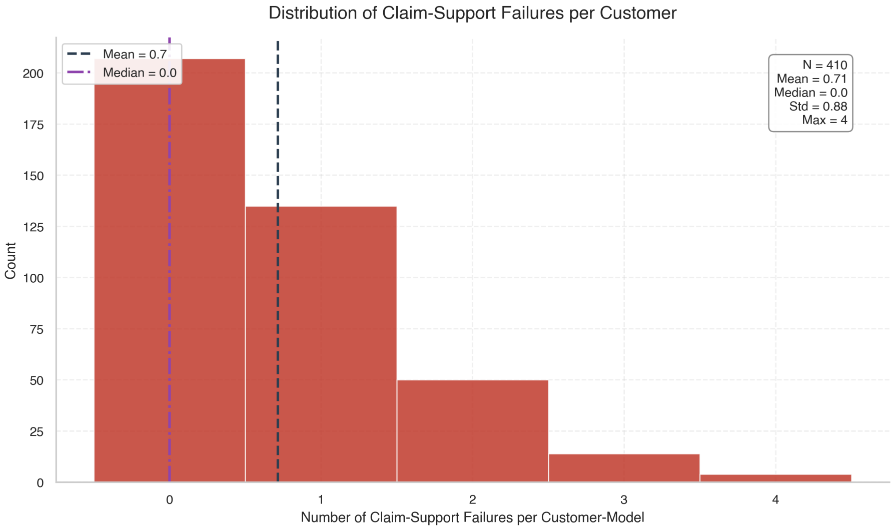


Figure 32: **Distribution of failed claim-support assertions (41-profile subset).** 207 of 410 customer-model pairs (50%) had zero claim failures.
